# Galectin-3 drives tau-associated neuroinflammation, white matter degeneration and proteomic dysregulation

**DOI:** 10.64898/2026.07.07.736964

**Authors:** Lluís Camprubí-Ferrer, Marzia Dell’Eva, Jesús Soldán-Hidalgo, Alberto Lerma-Aguilera, Laura R. Rodríguez, Javier Frontiñán-Rubio, Katryna Pampuščenko, Emil Axell, Erika Velasquez, Yiyi Yang, Henrik Ahlenius, Juan García-Revilla, Javier Vitorica, Antonio Boza-Serrano, José Luis Venero, Tomas Deierborg

**Affiliations:** Experimental Neuroinflammation Laboratory, Lund University, Sweden; Instituto de Biomedicina de Sevilla, IBiS/Hospital Universitario Virgen del Rocío/CSIC/Universidad de Sevilla, Spain; Stem cells, Aging and Neurodegeneration group, Lund University, Sweden; Oxidative Stress and Neurodegeneration Group, Universidad de Castilla-La Mancha, Ciudad Real, Spain; Biochemistry and Structural Biology, Department of Chemistry, Lund University, Lund 223 62, Sweden; Department of Biochemical Engineering, Lund University, Sweden

**Keywords:** tauopathy, galectin-3, neuroinflammation, microglia, white matter, myelin

## Abstract

Tau pathology is a central driver of neurodegeneration, yet the molecular mechanisms linking tau accumulation to neuroinflammation, metabolic failure, and white matter degeneration remain incompletely understood. Galectin-3 (Gal3) is an inflammation-associated lectin expressed by activated microglia and has been implicated in neurodegenerative disease progression. Here, we investigated whether Gal3 modulates tau-driven pathology across cellular, molecular, and systems levels.

Using the P301S tauopathy mouse model with genetic deletion of Gal3, we show that Gal3 loss robustly attenuates tau pathology across vulnerable brain regions, including cortex, hippocampus, and piriform–entorhinal cortex. Gal3 deletion reduced hyperphosphorylated and pathological tau species, normalized tau kinase signaling, and restored mitochondrial and vesicular trafficking pathways disrupted by tau accumulation. Proteomic and phosphoproteomic analyses revealed widespread normalization of tau-associated immune, metabolic, and trafficking pathways, with Tau-Gal3KO mice clustering closely with wild-type controls. In parallel, Gal3 deletion markedly reduced microglial activation and Gal3-positive inflammatory signatures, preserved white matter integrity, prevented axonal degeneration, and normalized oligodendrocyte and myelin abnormalities.

Functionally, Gal3 deficiency enhanced microglial myelin phagocytosis and lysosomal degradation both in vitro and in vivo, suggesting improved clearance of myelin debris under inflammatory stress. Cell-type–specific analyses further revealed restoration of mitochondrial complex I subunit expression in both excitatory neurons and parvalbumin-positive interneurons. Importantly, translational studies in human iPSC-derived neurons demonstrated that extracellular Gal3 exacerbates tau hyperphosphorylation and aggregation following tau seeding, effects that were reversed by pharmacological Gal3 inhibition.

Together, these findings identify Galectin-3 as a central upstream regulator linking tau pathology to neuroinflammation, proteomic dysregulation, mitochondrial dysfunction, and white matter degeneration. Targeting Gal3 represents a promising therapeutic strategy to mitigate tau-driven neurodegenerative processes.

## Introduction

Neurodegenerative diseases (NDs), such as Alzheimer’s disease (AD) and Frontotemporal Dementia (FTD), represent a growing and severe burden on global public health, characterized by progressive cognitive decline and neuronal loss (1). FTD is recognized as the second most common form of early-onset dementia (2). The neuropathology of FTD is heterogeneous, with a significant proportion of cases defined by the abnormal accumulation of the microtubule-associated protein Tau (FTD-tau) (3, 4). Given the absence of effective therapeutic or curative agents, investigating the core mechanisms driving FTD progression is critical for developing future interventions (5).

There is overwhelming evidence that neuroinflammation is a major factor contributing to the pathogenesis and accelerated progression of FTD (3, 6). This inflammatory state involves the chronic activation of the central nervous system’s innate immune cells, primarily microglia (6–8). Microglial activation is a pathological hallmark, detectable in vivo using PET imaging ligands in the frontotemporal regions, often preceding marked anatomical atrophy (9–11). Chronic microglial activation creates a sustained proinflammatory environment in the brain, which is central to disease development and progression (6). This neuroinflammatory setting is marked by dysregulated pro- and anti-inflammatory factors, including cytokines like TNF-α and IL-1β, which are linked to neuronal damage (5, 12).

A crucial element of FTD pathology is the compromise of white matter (WM) integrity (13). WM injury, involving the disruption of myelin and oligodendrocytes, is a pervasive phenomenon across neurodegenerative diseases, frequently appearing before the onset of clinical symptoms (13, 14). In FTD, diffusion tensor imaging (DTI) studies reveal a significant longitudinal decline in WM microstructural integrity, particularly in critical tracts like the corpus callosum (15, 16). Histological studies confirm that microglial activation is often more prominent in the frontal subcortical white matter of FTD patients compared to gray matter (17, 18). This suggests that the microglial response is localized and specialized within different brain regions (18). Indeed, microglial populations involved in WM homeostasis, often termed White Matter-Associated Microglia (WAMs), are essential for monitoring myelin health and clearing debris (19, 20). WM damage is exacerbated by neuroinflammation, where T cells and proinflammatory cytokines can directly damage oligodendrocytes (13, 21). Microglial activation in tauopathy specifically drives changes in microglial subtypes, increasing disease-associated microglia (DAM) at the expense of WAMs, potentially impairing the clearance of myelin debris and accelerating WM degeneration (19, 22).

Galectin-3 (Gal3), encoded by the LGALS3 gene, is a master regulator of microglial activation and inflammation (23, 24). It is predominantly expressed and released by reactive microglial cells in the compromised central nervous system (25, 26). Gal3 functions as an endogenous ligand for key receptors, including the Triggering Receptor Expressed on Myeloid cells 2 (TREM2) and Toll-like Receptor 4 (TLR4), thereby mediating or amplifying the microglial inflammatory response (24, 25). The expression of Gal3 is intricately linked to the neurodegenerative microglial phenotype (MGnD/DAM), which is often driven by the TREM2-APOE pathway (7, 27). In tauopathies, Gal3 is strongly associated with tau aggregates in brain tissue and exacerbates microglial activation and tau transmission (26, 28). Experimental evidence suggests a detrimental role for Gal3, as its inhibition reduces proinflammatory cytokines and alleviates pathology in models of neurodegeneration (24, 28).

In FTD, Gal3 levels are significantly upregulated in brain tissue, CSF, and serum compared to controls, underlining the crucial role of Gal3-expressing microglia in the neurodegenerative mechanism (29). This upregulation demonstrates a remarkable neuropathological specificity, being significantly higher in FTD cases with Tau pathology (FTD-tau), particularly in MAPT mutation carriers (29). This finding is important because MAPT mutation carriers often display strong microglial activation (18, 30) and the elevation of Gal3 in CSF correlates with markers of neuronal injury, such as total tau (t-tau) (29). The finding that microglial activation, often indicated by markers like Gal3, may be an early event in MAPT mutation carriers suggests new possibilities for early anti-neuroinflammatory treatments (29, 31).

The objective of the current study is to characterize the specific and detrimental role of Galectin-3 in the pathogenesis of FTD, focusing on its link to Tau pathology and its expression within the specialized inflammatory landscape of the FTD brain, specially on the impact of the pathology on white matter integrity along with the underlying mechanism driving microglial activation.

## Materials and methods

### Animals

All animal experiments were conducted in accordance with Spanish and European Union regulations (RD53/2013 and Directive 2010/63/EU) and were approved by the Animal Research Committees of the University of Seville and the Malmö/Lund Animal Ethics Committee (permit numbers M30-16 and Dnr 5.8.18-01107/2018). Male and female 9-month-old mice were used in this study, including P301S tau transgenic mice (hereafter referred to as Tau; Jackson Laboratory), galectin-3 knockout mice (Gal3KO; originally obtained from Dr. K. Sävman, University of Gothenburg), and wild-type littermates (WT). All mouse lines were maintained on a C57BL/6 background. The P301S mice express human four-repeat tau with one N-terminal insert carrying the P301S mutation under the control of the mouse prion protein (Prnp) promoter (B6;C3-Tg(Prnp-MAPT*P301S)PS19Vle/J). Animals were housed under controlled environmental conditions at a constant temperature of 22 °C, with ad libitum access to food and water, and maintained on a 12-h light/12-h dark cycle.

### Tissue Collection and Processing

Mice were deeply anesthetized with isoflurane (4% in oxygen) and transcardially perfused with ice-cold phosphate-buffered saline (PBS; Gibco). Following perfusion, brains were rapidly removed and hemisected, with each hemisphere processed for distinct downstream applications.

One hemisphere was fixed in 4% paraformaldehyde (PFA; Histolab) for 24 h, then cryoprotected in PBS containing 30% sucrose (Millipore) and 0.01% sodium azide (Sigma-Aldrich) at 4 °C until sectioning. Fixed tissue was sectioned coronally at 35 μm using a microtome (Leica SM2500 DR), collected into six serial sets, and stored at −20 °C in cryoprotectant solution (30% sucrose, 30% ethylene glycol, and 40% phosphate buffer) until further use.

From the contralateral hemisphere, the prefrontal cortex (PFC), hippocampus (HC), and piriform/entorhinal cortex (Pir-Ent) were dissected at the time of sacrifice, snap-frozen on dry ice, and stored at −80 °C. PFC tissue was homogenized on ice using glass Dounce homogenizers, and sequential protein extraction was performed based on solubility in three buffers applied in order: PBS (Gibco), radioimmunoprecipitation assay buffer (RIPA; Fisher Scientific), and 4% sodium dodecyl sulfate (SDS; Sigma-Aldrich) with 8 M urea (GE Healthcare). For each extraction step, tissue was resuspended at a ratio of 10 μL buffer per mg of original tissue, centrifuged at 20,000 × g for 40 min at 4 °C, and the supernatant collected. The resulting pellet was subsequently used for the next extraction step. All protein fractions were stored at −80 °C until analysis. HC and Pir-Ent samples were directly homogenized in RIPA buffer for proteomic analyses, and their RIPA-insoluble fractions were further solubilized using 4% SDS-8 M urea.

### Immunohistochemistry

Two anatomically distinct coronal levels were analyzed: bregma −1.94 mm (referred as Aanterior) and bregma −3.08 mm (referred as Posterior). Immunohistochemical staining was performed using a free-floating 3,3′-diaminobenzidine (DAB) method.

Sections were washed three times for 5 min under agitation in phosphate-buffered saline (PBS), followed by three washes in PBS containing 1% Triton X-100. Endogenous peroxidase activity was quenched by incubation in 99% methanol (Acros Organics) with 1% hydrogen peroxide for 15 min, after which sections were washed three times in PBS with 1% Triton X-100. Sections were then permeabilized in PBS with 1% Triton X-100 for 30 min and blocked for 60 min in blocking solution consisting of PBS, 1% Triton X-100, and 5% bovine serum albumin (BSA; Sigma-Aldrich). Sections were incubated overnight at 4 °C under gentle agitation with primary antibodies diluted in blocking solution. Antibodies and dilutions are listed in Table 1. Following overnight incubation, sections were equilibrated for 30 min at room temperature and washed three times for 5 min in PBS with 1% Triton X-100. Sections were then incubated with the appropriate secondary antibodies diluted in blocking solution for 1 h at room temperature under agitation (see Table 1 for details). After washing, sections were incubated with the VECTASTAIN Elite ABC reagent (Vector Laboratories) prepared according to the manufacturer’s instructions for 1 h at room temperature under agitation. Following additional washes, immunoreactivity was visualized using a DAB solution (Vector DAB kit; Vector Laboratories) prepared in distilled water. Sections were incubated for approximately 2 min, or until optimal signal-to-noise contrast was achieved, depending on the primary antibody. The reaction was stopped by washing sections three times in PBS. Sections were mounted onto Superfrost Plus microscope slides (Menzel-Gläser) and dehydrated through graded ethanol solutions (Solveco): 70% ethanol for 5 min, 90% ethanol for 3 min, 96% ethanol twice for 3 min each, and 100% ethanol for 3 min. Slides were then cleared in xylene (Sigma-Aldrich) twice for 6 min each, air-dried briefly, and coverslipped using Pertex mounting medium (Histolab). All brightfield images were taken with a Zeiss AxioImager microscope using an objective with a 20x magnification.

### Immunofluorescence

#### Brain slices

Brain slices were washed 3 times for 5 minutes and incubated for 1 hour in a blocking solution of PBS + 0.25% Tween20 + 10% Normal Donkey Serum (NDS, Millipore). After blocking, primary antibodies were added in PBS + 0.25% Tween20 + 10% NDS and an overnight incubation at 4°C under agitation was performed. A list of the antibodies used immunostainings can be found in Table 1. Slices were then washed 2 times for 5 minutes in PBS + 0.25% Tween20. Appropriate donkey-raised secondary Alexa Fluor antibodies (Thermo Fisher) were then added for 2 hours at room temperature in the dark. After secondary antibody incubation the slices were washed and DAPI (Thermo Fisher) was added for 5 min, washed with PBS and then mounted in Super Frost slides and coverslips were added with ProLong Glass Antifade mounting medium (Thermo Fisher). Confocal images were taken with Nikon A1 confocal microscope or Leica SP8 laser scanning confocal microscope.

#### Primary cell cultures

Cell cultures were fixed with 4% PFA for 15 min, followed by 3 washes with PBS for 5 min. To quench free aldehyde groups, cells were incubated with 0.1 M glycine for 10 min. Following quenching, cells were permeabilized with 0.1% saponin for 10 min. and blocked with 15% bovine serum albumin (BSA) for 30 min. at room temperature. Cell cultures were incubated overnight at 4 °C with primary antibody for MAP2 (Table 1), and washed 3 times with PBS for 5 min. Then, secondary donkey-raised antibody in 5% BSA was added for 2h at room temperature. After 3 washed with PBS for 5 min, microglial cells were labeled with 1 µg/ml isolectin GS-IB_4_ from *Griffonia simplicifolia*, Alexa Fluor 488 conjugate (Thermo Fisher) for 20 min at 37 °C. Additionally, 1µg/ml DAPI was added to label cell nuclei (Thermo Scientific). Cell cultures were imaged under an epifluorescence Olympus IX70 microscope in five randomly chosen microscopic fields at 20x magnification. Manual quantification of neurons and microglia was carried out with ImageJ (version 1.8.0). Neuronal and microglial cell numbers in control group were normalized to 100%, and changes in tau or/and galectin-3 protein treated groups were expressed relative to control.

#### IPS-derived neurons

Cells were fixed in 4% PFA for 15 min, washed in PBS, blocked and permeabilized for 1 h in PBS containing 0.1% Triton X-100 and 5% normal donkey serum, and incubated overnight at 4 °C with primary antibodies. After washing, sections were incubated for 1 h at room temperature with secondary antibodies and DAPI in blocking solution.

#### ELISA

Galectin-3 levels were measured in RIPA-soluble brain fractions utilizing the DuoSet ELISA - Mouse Galectin-3 (R&D Systems), following manufacturer’s instructions.

#### Mesoscale Discovery Platform

Levels of total tau (totTau) and phosphorylated tau (pTau) were quantified in the SDS-urea-insoluble protein fraction of HC samples. Total tau and pTau phosphorylated at Thr231 were measured using the MSD Multi-Spot Phospho (Thr231)/Total Tau assay (Meso Scale Discovery). The assay detection ranges were 0.55-1000 ng/mL for total tau and 6.6-636 ng/mL for pTau (Thr231). Phosphorylated tau species at Thr181 (pTau181) and Thr217 (pTau217) were measured in the same SDS-urea-insoluble fractions from HC using the S-PLEX Human Tau (pT181) and S-PLEX Human Tau (pT217) kits (Meso Scale Discovery), respectively. Detection ranges were 0.00012-2.21 ng/mL for pTau181 and 0.00095-3.63 ng/mL for pTau217. All assays were performed according to the manufacturers’ instructions. All the plates were read by a QuickPlex Q120 according to the manufacturer’s instructions. The data were analyzed with MSD Discovery Workbench software. Some of the samples were under the lowest detection limit and were excluded from the final statistical analysis.

#### Transmission electron microscopy

Corpus callosum and corticospinal tract regions were isolated from 4% PFA fixed brains. The corpus callosum was isolated using a scalpel, and the corticospinal tract utilizing a 1 mm disposable biopsy punch with plunger (Miltex GmbH). Samples were initially fixed in 2% paraformaldehyde and 2% glutaraldehyde (Ted Pella; from a 25% stock solution) in phosphate-buffered saline (PBS) for 1 h at room temperature. Following fixation, samples were rinsed thoroughly in PBS (five washes of 10 min each).

Post-fixation was performed in 1% osmium tetroxide (OsO□; Ted Pella; prepared from 0.5-2% stock solutions) for 1 h, followed by rinsing in PBS. Samples were then dehydrated through a graded acetone series consisting of 30%, 50%, 70%, 80%, and 90% acetone for 5-10 min each, followed by three changes of 100% acetone for 10 min each.

Resin infiltration was carried out by incubating samples overnight in a 1:1 mixture of acetone and Epon resin with gentle agitation, followed by incubation in pure Epon for 4-5 h. Samples were subsequently transferred into embedding molds containing fresh Epon resin, oriented appropriately, and polymerized at 60 °C for 48 h.

Embedded samples were trimmed and ultrathin sections (60 nm) were cut and mounted on grids. Sections were contrasted with 4% uranyl acetate for 20 min at 40 °C and 2% lead citrate for 1-2 min at room temperature. Imaging was performed using a Thermo Fisher Talos L120C transmission electron microscope operated at 120 kV.

#### Recombinant Tau preparation

Tau protein preparation and aggregation were performed as previously described (32). For the present study, tau protein comprised the paired helical filament-enriched region of human tau441 (residues 304–380). Tau fibrils were generated by incubating the protein at 37 °C for 24 h under constant stirring in LoBind tubes (Eppendorf) containing a Teflon-coated stir bar. Complete fibril formation was confirmed using a Thioflavin T (ThT) fluorescence assay. Tau oligomers were collected at the half-maximal aggregation time point (t½) of the ThT kinetics, prepared in parallel without the addition of ThT dye.

### Cell cultures

#### BV2 cell cultures

BV2 WT and BV2 Gal3KO cells were produced and maintained as previously published (33). Briefly, cell media consisted of Dulbecco’s Modified Eagle Medium GlutaMax (DMEM GlutaMax, Gibco) supplemented with 10% heat inactivated Fetal Bovine Serum (FBS, Gibco) and 1% penicillin-streptomycin (Cytiva) and kept at 37°C 5% CO_2_. For myelin and Tau treatments, media was exchanged to FBS-free media. For live-cell imaging experiments, Hoechst, LysoTracker and MitoTracker (Invitrogen) were used to stain nuclei, lysosomes and mitochondria respectively. Live-cell images were taken under 37°C 5% CO_2_ in Operetta CLS High Content Screening (Revvity) by spinning disk technology with a 40x water immersion objective. For myelin addition to cell cultures, purified myelin was isolated from WT mice as previously published (34). Myelin was preincubated for 1h at 37°C with Fluoromyelin Red (Thermo Fisher; 2 parts of myelin to 1 part of Fluoromyelin), and added to cultures at 1:150.

Tau fibrils were cup-horn sonicated prior to addition to cell cultures, in a Q700 sonicator (Qsonica) in cycles of 1 sec on and 1 sec off, with an amplitude of 70% for 6 minutes in total. Tau fibrils were preincubated for 30 min at 37°C with Amytracker 630 (Ebba Biotech; 9 parts of tau to 1 part of Amytracker), and added to cultures at 0.1 µM.

#### Primary microglial cell cultures

Primary neuronal-glial co-cultures were prepared from cerebella collected from WT and Gal3KO postnatal mice on 5-7 day. Tissue was immediately transferred to ice-cold HBSS supplemented with 1 % penicillin-streptomycin. After meninges and blood vessels were removed, chopped tissue was transferred to Versene solution and incubated in 37 °C water bath for 5 min. Tissue was triturated and then centrifugation at 300 × g for 5 min. Supernatant was discarded, and cell pellet was resuspended in cell growth medium consisting of DMEM Glutamax supplemented with 5% FBS, 5 % horse serum, 13 mM glucose, 20 mM KCl and 1 % penicillin-streptomycin. Cell suspension was passed through 40 µm nylon mesh strainer and subsequently plated at a density of 200,000 cells per well in 48-well plates pre-coated with 0.001% poly-L-lysine solution. Cell cultures were maintained at 37 °C in a humidified atmosphere containing 5% CO□ for 7-8 days before treatments. Treatments included: 1) single tau treatment (200 nM) for 72h; 2) sequential tau and Gal3 treatment, in which tau (200 nM) was added for 24h after which Gal3 (1 µM) was added for 48h; 3) co-aggregation tau-Gal3 treatment, in which tau was preincubated with Gal3 at 1:0.1 or 1:2 ratios and added for 72h, preincubation was performed for 3 h at 22 °C with agitation at 14,000 rpm in LoBind tubes (Eppendorf). As controls, Gal3 was added alone at concentrations of 20 nM (for 1:0.1 experiments), 400 nM (for 1:2 experiments) or 1µM (for sequential experiments).

Tau fibrils were sonicated prior to addition to the culture, as previously described.

### Human iPSC lines, generation of iPSC-derived neurons and culture conditions

The human male wild-type induced pluripotent stem cell (iPSC) line was obtained from the NINDS Human Cell and Data Repository (NH50191). An isogenic MAPT-mutant iPSC line was generated in the same genetic background by introducing the homozygous P301L point mutation in the MAPT gene. Genome editing was performed using CRISPR/Cas9-mediated homologous recombination, employing the single-stranded oligodeoxynucleotide (ssODN) repair template (GTCCAGTCCAAGTGTGGCTCAAAGGATAATATCAAACACGTCCTGGGAGGCGGCAGTGTGAGTACCTTCACACGTCCCAT) and the guide RNA (gRNA) sequence GATAATATCAAACACGTCCC.

Comprehensive quality control analyses were performed for both the parental and edited iPSC lines, including assessment of cellular morphology, verification of the undifferentiated state, Sanger sequencing of the edited genomic region, copy number variation analysis, off-target evaluation, cell line authentication, and molecular karyotyping.

iPSC lines were cultured on Matrigel hESC-Qualified Matrix-coated plates (Corning,) and maintained in Essential 8 medium (eTeSR; STEMCELL Technologies). Cells were passaged upon reaching 80-90% confluency using Accutase cell dissociation reagent (Gibco) in the presence of 10 μM ROCK inhibitor Y-27632 (Selleckchem). The ROCK inhibitor was removed 24 hours after passaging. Cells were maintained at 37°C in a humidified incubator with 5% CO□, and maintenance medium was replaced every other day. Cultures were routinely tested for mycoplasma contamination.

Transcription factor forward programming protocols were used to convert human WT and MAPT iPSCs into induced cortical neurons (iNs). Transcription factors (TFs) were delivered as lentiviral vectors under the control of a TetO-response promotor. TF gene expression was induced by co-transduction of the gene of interest along with the reverse tetracycline-controlled transactivator (rtTA), which was expressed under the constitutive ubiquitin promotor (35). Doxycycline (2.5μg/mL, Sigma-Aldrich) was added to the culture media daily to activate rtTA and induce transgene expression via the Tet-On system.

For generation of iNs, 4.3×105 iPSCs were plated per well of 6-well culture plates (Corning) using Growth Factor Reduced (GFR) Basement Membrane Matrix (Corning) and eTeSR supplemented with 10μM Ri (day-2). On day-1, iPSCs were transduced with lentiviral vectors containing plasmids for the TF Ngn2 and rtTA. After overnight incubation at 37°C, media of the infected cells was removed and differentiation was induced by adding fresh eTeSR containing doxycycline (day 0). On day 1, media was changed to BrainPhys Neuronal Medium (BPm, STEMCELL Technologies) with doxycycline and puromycin (2.5μg/mL, Gibco) was added for selection. BPm was supplemented with 2% B27 (Gibco) and 1% N2 supplement (Gibco). Full media change was performed on day 2 and 3 with the same media composition. On day 4, media was changed using BPm and doxycycline. Complete media change was conducted on day 5 and 6 using BPm supplemented with doxycycline and the neural factors human neurotrophin-3 (NT3, 10ng/mL, Peprotech) and brain-derived neurotrophic factor (BDNF, 10ng/mL, Peprotech). On day 7, cells were detached using accutase and replated in GFR coated wells containing BPm supplemented with doxycycline, NT3 and BDNF. 240000 neurons were seeded on 24-well format (ibidi). After replating for iN mono-culture, half-media change was performed on day 8 using BPm supplemented with doxycycline, NT3 and BDNF. From day 9 onwards, half media change was performed every 2-3 days. Treatments were administered according to the experimental timeline as follows. Pre-sonicated tau fibrils were applied on day 9, prepared as described above. Recombinant extracellular galectin-3 (Gal3; (33)) was administered on day 12. The galectin-3 inhibitor TD139 (kindly provided by Ulf Nilsson) was applied on day 12 or day 13, as indicated. At day 14, cells were rinsed once with sterile PBS and fixed with 4% PFA.

### Image analysis

Unless indicated otherwise, all images were analyzed using Fiji ImageJ version 2.13.1. For MBP content within Iba1 cells, Microscopy Image Analysis Software IMARIS (Oxford Instruments) was used.

### Proteomic and phosphoproteomic analysis

#### Sample preparation

Protein digestion (50 µg of proteins per sample) was performed using the S-Trap™ 96-well plate method (ProtiFi) according to the manufacturer’s instuctions. In brief, tissue lysates were diluted to 45 uL with a buffer containing dithiothreitol (DTT) and sodium dodecyl sulphate (SDS) to a final concentration of 15 mM DTT and 5 % SDS, and incubated at 95 °C for 10 min. The proteins were then alkylated with 50 mM iodoacetamide (IAA) for 30min at room temperature in the dark. Then, 350 µL of S-Trap binding buffer (90% Methanol, 100 mM TEAB) was added and samples were transferred to the S-Trap 96-well digestion plate, which was centrifuged for 2 min at 2,000 x g. Captured proteins were washed 3 times with 200 µL of S-Trap binding buffer using brief centrifugations (2 min at 1,000 × g).Finally, 125 µL of digestion buffer (50 mM TEAB) containing trypsin (sequencing grade modified trypsin, Promega Biotech AB) at 1:50 enzyme-to-protein ratio was added on top of the filters, incubating overnight at 37 °C. On the following day, the peptides were eluted in three steps, first with 80 µL of 50 mM TEAB, then with 80 µL of 0.2% formic acid, and finally with 80 µL of 50% acetonitrile (ACN) containing 0.2% formic acid. All peptides were dried down in a vacuum concentrator and for the MS analysis they were resuspended in 2% ACN containing 0.1% trifluoroacetic.

Peptide concentrations were determined using the Pierce Quantitative Colorimetric Peptide Assay. From each sample, 1 µg of peptides was injected into the nLC-MS/MS system for global proteomic analysis. The remaining peptide digest was subjected to C18-based cleanup using the automated AssayMAP Bravo system (Agilent Technologies) for subsequent phosphoproteomic analysis. Following desalting, phosphopeptides were enriched by Fe(III)-IMAC affinity chromatography as described previously (36). The enriched fractions were then resuspended in 2% acetonitrile (ACN) containing 0.1% trifluoroacetic acid (TFA) and analyzed by nLC-MS/MS.

#### nanoLC-MS/MS analysis for proteomics

The nLC-MS/MS analysis was performed on an Exploris 480 mass spectrometer coupled to a Vanquish Neo nano UPLC system (Thermo Scientific), with an EASY-Spray ion source. Samples were run in a randomized order using high resolution data independent acquisition (HR-DIA). All samples were loaded onto an Acclaim PepMap 100 C18 (75 µm × 2 cm, 3 µm, 100 Å, nanoViper) trap column and separated on an Acclaim PepMap RSLC C18 column (75 µm × 50 cm, 2 µm, 100 Å) (Thermo Scientific) using a flow rate of 300 nL/min, a column temperature of 60°C. A 90 min non-linear gradient was applied for separation, using solvents A (0.1% formic acid) and B (0.1% formic acid in 80% ACN), increasing solvent B from 5 to 25% in 75 min, then to 32% in the next 9 min, and to 45% in 6 min. Finally, the gradient increased to 95% solvent B in 2 min, continuing for another 8 min. In the HR-DIA analysis, a complete acquisition cycle consisted of 3 MS1 full scans, each followed by 18 MS2 DIA scans with variable isolation windows. Full scans were acquired at m/z 375-1455, with a resolution of 120,000 (at 200 m/z), target AGC value of 300% and maximum injection time of 45 ms. The MS2 scans were acquired with a resolution of 30,000, fragmentation with NCEs of 27, 30 and 32, target AGC value of 1000%, automatic maximum injection time, fixed first mass of 200m/z, and 1 m/z overlap between isolation windows). Isolation windows were 13.0, 16.0, 26.0 and 61.0 m/z with loop counts of 27, 13, 8, 6, respectively.

#### nanoLC-MS/MS analysis for phosphoproteomics

The nLC-MS/MS analysis was performed on an Exploris 480 mass spectrometer coupled to a Vanquish Neo nano UPLC system (Thermo Scientific), with an EASY-Spray ion source. Phosphopeptide samples were analyzed using data-dependent acquisition (DDA) with a top 15 method. All samples were loaded onto an Acclaim PepMap 100 C18 trap column (75 μm × 2 cm, 3 μm, 100 Å, nanoViper) and separated on an EASY-Spray RSLC C18 analytical column (75 μm × 25 cm, 2 μm, 100 Å) using a flow rate of 300 nL/min and a column temperature of 45 °C. A nonlinear gradient was applied for separation using solvents A (0.1% formic acid) and B (0.1% formic acid in 80% acetonitrile), increasing solvent B from 5 to 25% over 110 min, then to 32% in the next 10 min, and to 45% in 8 min. Finally, the gradient increased to 95% solvent B in 2 min and was maintained for an additional 13 min. Full MS1 scans were acquired at m/z 350-1750 with a resolution of 120,000 (at m/z 200), a target AGC value of 300%, and a maximum injection time of 45 ms. The 15 most intense precursor ions were selected for higher-energy collisional dissociation (HCD) using a normalized collision energy of 27. MS2 scans were acquired at a resolution of 45,000 with a target AGC value of 100%, a maximum injection time of 120 ms, and a dynamic exclusion of 30 s.

#### Database search for proteomic data

MS files were analyzed using Spectronaut vs18.4 (Biognosys AG) in library-free mode, using directDIA factory default settings and MS1 quantification. The UniProt mouse protein database (17,184 entries) extended with human TAU was used for the database search. Carbamidomethylation on cysteine and oxidation on methionine residues were considered as fixed and variable modifications, respectively. A maximum of two missed cleavages were allowed. A 1% FDR was applied at both peptide and protein levels.

#### Database search for phosphoproteomic data

MS data were analyzed using Proteome Discoverer v2.5 (Thermo Scientific) with the SEQUEST HT search engine. The UniProt mouse protein database (17,184 entries) extended with human TAU was used for the database search. Trypsin was selected as the protease, allowing up to two missed cleavages. The precursor and fragment mass tolerances were set to 10 ppm and 0.02 Da, respectively. Carbamidomethylation of cysteine was set as a fixed modification, while methionine oxidation, phosphorylation on serine, threonine, and tyrosine, and protein N-terminal acetylation were defined as variable modifications. Phosphorylation site localization was evaluated using ptmRS with a site probability threshold >75%. A 1% FDR was applied at both peptide and protein levels.

### Bioinformatic analysis

Proteomic and phosphoproteomic analyses were performed independently for hippocampus and entorhinal cortex. LFQ intensities were normalized using variance-stabilizing normalization (vsn). Proteins were retained if they contained at least three valid measurements per experimental group. Differential abundance was assessed using the limma package in R, fitting a linear model without intercept and including sex as a covariate. Adjusted p-values were computed using the Benjamini-Hochberg (BH) procedure. Proteins and phosphosites with adj. p-values ≤ 0.05 were considered differentially abundant.

Visualizations included volcano plots generated from log2 fold changes and adjusted p-values, and heatmaps constructed from vsn-normalized abundances of differentially abundant proteins, scaled by gene-wise z-scores. Venn diagrams were used to illustrate shared differentially abundant proteins across comparisons.

#### Pathway Enrichment Analysis

Gene set enrichment analyses (GSEA) were conducted for KEGG and Reactome pathways. Protein identifiers were mapped to ENTREZ gene IDs using the AnnotationDbi (v1.68.0) package in R, and only proteins included in the limma analysis were retained in the ranked background. Enrichment was performed using GSEA, employing a composite ranking metric defined as score=sign(log2FC)×log10(FDR).

Ranked gene lists were analyzed using clusterProfiler (v4.14.0) package with minimum and maximum gene set size of 10 and 500, with BH procedure for multiple testing corrections. KEGG pathway visualizations illustrating differential abundance patterns across comparisons were generated using the pathview (v1.46.0) package in R. Shared significant pathways in at least TvsTG3KO and TvsWT comparisons were included in the heatmap along with its normalized enrichment score (NES). Analyses were performed in R version 4.4.3.

### Statistical analysis

Statistical analyses were performed using GraphPad Prism version 10.0.0. All data sets were subjected to the ROUT outlier test with a Q of 1% and all identified outliers were eliminated from the analysis. The type of statistical test was chosen specifically for each graph, based on whether the data was normally distributed or not and whether the variances between groups were comparable or not. Normality was checked by performing a Normality test and differences in variances were evaluated through an F-test, both performed on GraphPad Prism. Normally distributed data sets with equal variances were subjected to an unpaired t-test, while normally distributed data sets with different variances were subjected to a Welch test. All not-normally distributed data sets were subjected to a Mann-Whitney test. A significance threshold of p < 0.05 was applied. Exact p-values, including those approaching significance, are reported in the figure panels or legends. The specific statistical tests used for each dataset are indicated in the corresponding figure legends.

## Results

### Galectin-3 deletion attenuates tau hyperphosphorylation and pathological tau accumulation across vulnerable brain regions

The investigation into the consequences of deleting Gal3 in a tauopathy mouse model (TauGal3KO versus Tau) directly addresses the hypothesis that Gal3, widely recognized as a marker of detrimental microglia phenotypes in neurodegenerative disorders like Alzheimer’s disease (AD) and frontotemporal dementia (FTD), exacerbates core pathology. Gal3 is highly upregulated in the brains of AD patients and associated with microglia surrounding amyloid plaques, acting as a ligand for critical immune receptors such as Toll-like Receptor 4 (TLR4) and Triggering Receptor Expressed on Myeloid Cells 2 (TREM2) (24, 25), thereby driving proinflammatory signaling. Our detailed analysis demonstrates that the absence of Gal3 significantly reduces the pathological burden in regions vulnerable to Tau aggregation. First, we assessed the hyperphosphorylated of Tau (AT8^+^ cells/mm^2^) in isocortex as well as in the piriform-entorhinal cortex (PirEnt) in male and female mice (Fig. 1 A-D), in both cases we analyzed the anterior and posterior areas. Notably, only Tau female mice lacking Gal3 in revealed reduced pathological AT8 load mice compared to the Tau mice. This reduction was noted in the anterior and posterior sections in both areas, isocortex (p=0045; p=0,028. Fig. 1A-B) and PirEnt (p=0,036, p=0,022. Fig. 1C-D), confirming an ameliorating effect of Gal3 deletion on Tau accumulation in cortical areas.

**Fig. 1.**
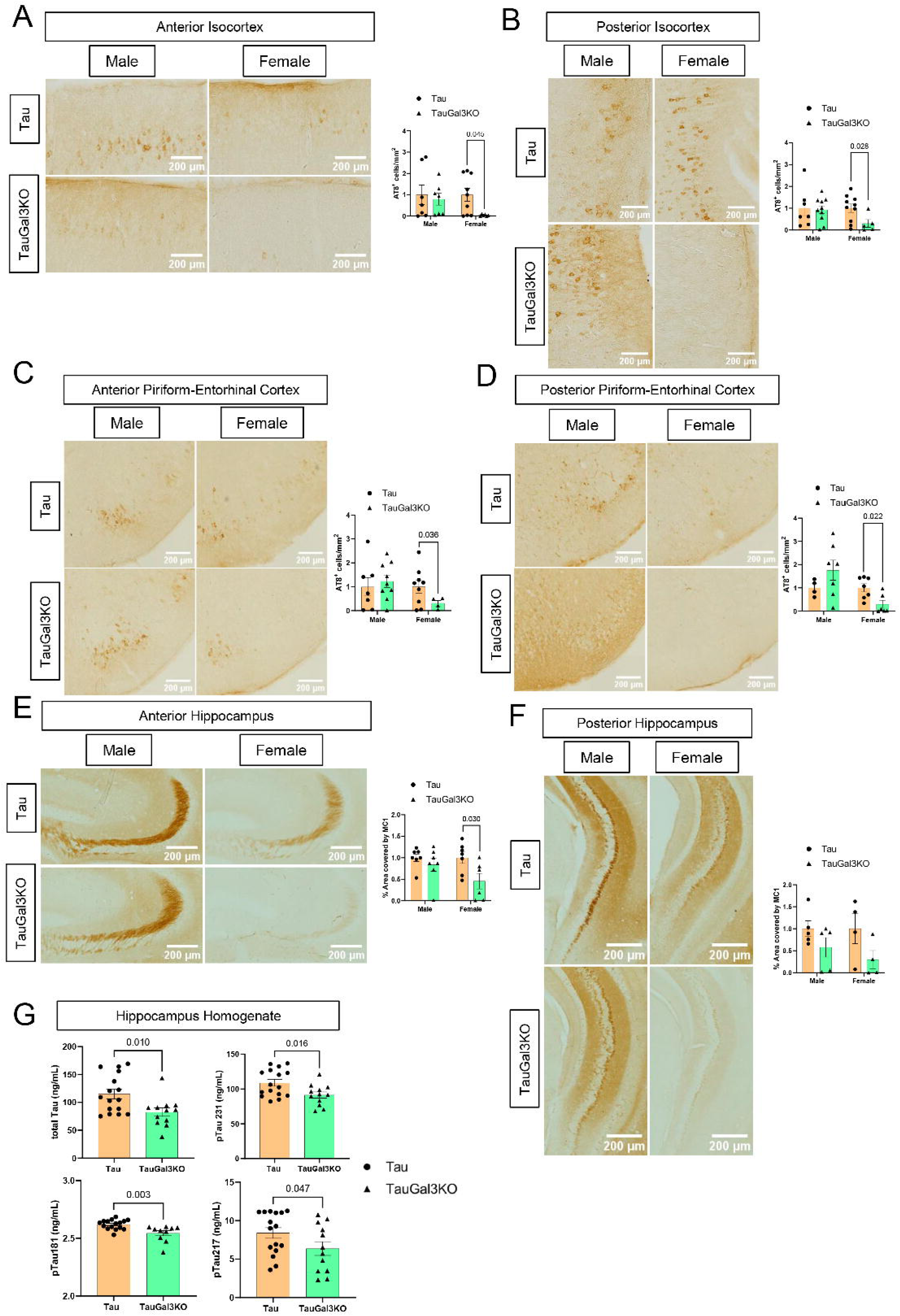
Galectin-3 deletion reduces hyperphosphorylated and pathological tau burden across cortical and hippocampal regions. **(A)** Representative images of 9-month-old male and female Tau and TauGal3KO mice anterior Isocortex (specifically the CA3 region) stained with AT8 and quantification of the number of AT8-positive (AT8^+^) cells per mm^2^. (**B)** Representative images of posterior Isocortex and quantification of AT8-positive (AT8^+^) cells per mm^2^. **(C)** Representative images of anterior Piriform-Entorhinal cortex and quantification of AT8-positive (AT8^+^) cells per mm^2^. **(D)** Representative images of posterior Piriform-Entorhinal cortex and quantification of AT8-positive (AT8^+^) cells per mm^2^. **(E)** Representative images of anterior hippocampus stained with MC1 and quantification of the percentage of area covered by MC1. **(F)** Representative images of posterior hippocampus stained with MC1 and quantification of the percentage of area covered by MC1. **(G)** Concentration of total tau, pTau231, pTau181 and pTau217 in SDS-Urea-soluble hippocampi homogenates. TauGal3KO are normalized against same-sex Tau mice. All values are expressed as individual experimental replicates with mean ±SEM. Unpaired t-test was performed on all normal data sets with equal variances (A males, B, C, D males, E, F females), Welch’s test was performed on all normal data sets with different variances (A females), and Mann-Whitney test was performed on all non-normal data sets (D females, F males). Significant p-values are shown.

Moving to the hippocampus (HC), a critical structure highly susceptible to neurodegeneration (37), we quantified the percentage of area covered by the MC1 marker, which specifically labels the pathological conformation of Tau. Mice lacking Gal3 (TauGal3KO) showed a significantly lower MC1 area coverage compared to Tau mice in both the anterior and posterior hippocampus (p=0.030 anterior hippocampus. Fig 1E-F). The histological findings were strongly corroborated by biochemical analysis of the hippocampal homogenates (Fig. 1G). Specifically, levels of Total Tau were significantly reduced in TauGal3KO mice (p=0.016). The concentration of pTau231, a marker of early tau phosphorylation, was significantly lower in TauGal3KO mice compared to Tau mice (p=0.010). Other key hyperphosphorylated species, pTau217 (p=0.047) and pTau181 (p=0.003), also displayed statistically significant reductions in mice lacking Gal3. Significant reductions were also observed in the TauGal3KO group across multiple phosphorylation markers in hippocampus and the PirEnt, including pTau T202 (p=0.008), pTau T152 (p=0.009), and pTau S206 (p=0.002) (Suppl. Fig. 1A-B). Pathological sites involving multiple phosphorylation events also showed reductions, such as pTau S46 T50 (p=0.027) and pTau T202 S206 (p=0.039) in the hippocampus homogenate of TauGal3KO mice (Suppl. Fig. 1A-B)

In conclusion, these findings underscore that Gal3 plays a detrimental regulatory role in neurodegeneration. Its absence leads to a broad attenuation of core tau pathology markers, including reduced hyperphosphorylation, less pathological conformation, and lower biochemical loads of toxic tau species across vulnerable brain regions. This evidence supports the concept that Gal3 acts as a central upstream regulator driving the inflammatory environment that exacerbates tau pathogenesis.

### Galectin-3 drives tau-associated microglial activation and neuroinflammation

The analysis of microglial cells confirmed that the presence of tauopathy strongly induced a neuroinflammatory response that was characterized by the expression of Gal3, an established marker of activated microglia. Tau mice exhibited a robust increase in microglial load (Iba1 area coverage) across multiple vulnerable brain regions and white matter tracts, including the Corpus Callosum, Fimbria, and Corticospinal tract, compared to wild-type (WT) mice (p<0.001 in most regions. Fig. 2A-B). Critically, the deletion of Gal3 in TauGal3KO mice significantly attenuated this microglial activation, resulting in markedly lower Iba1 area coverage compared to Tau mice in nearly all regions analyzed (p<0.001 in most regions, e.g., Hippocampus/White Matter, Corpus Callosum. Fig. 2A-B). In Tau mice, this neuroinflammatory phenotype was characterized by high Gal3 expression. Quantitatively, the proportion of activated microglia (Iba1^+^) that co-expressed Gal3 (Gal3^+^/Iba1^+^) was significantly elevated in Tau mice compared to WT in the Corticospinal tract, Fimbria and Corpus Callosum (all p≤0.001, Fig. 2C-D; Suppl. Fig. 2A-B). This increase was confirmed biochemically, with Gal3 protein levels in tissue homogenates showing significant upregulation in the hippocampus (p=0.004) and prefrontal cortex (p=0.045) of Tau mice compared to WT (Fig. 2E).

**Fig. 2.**
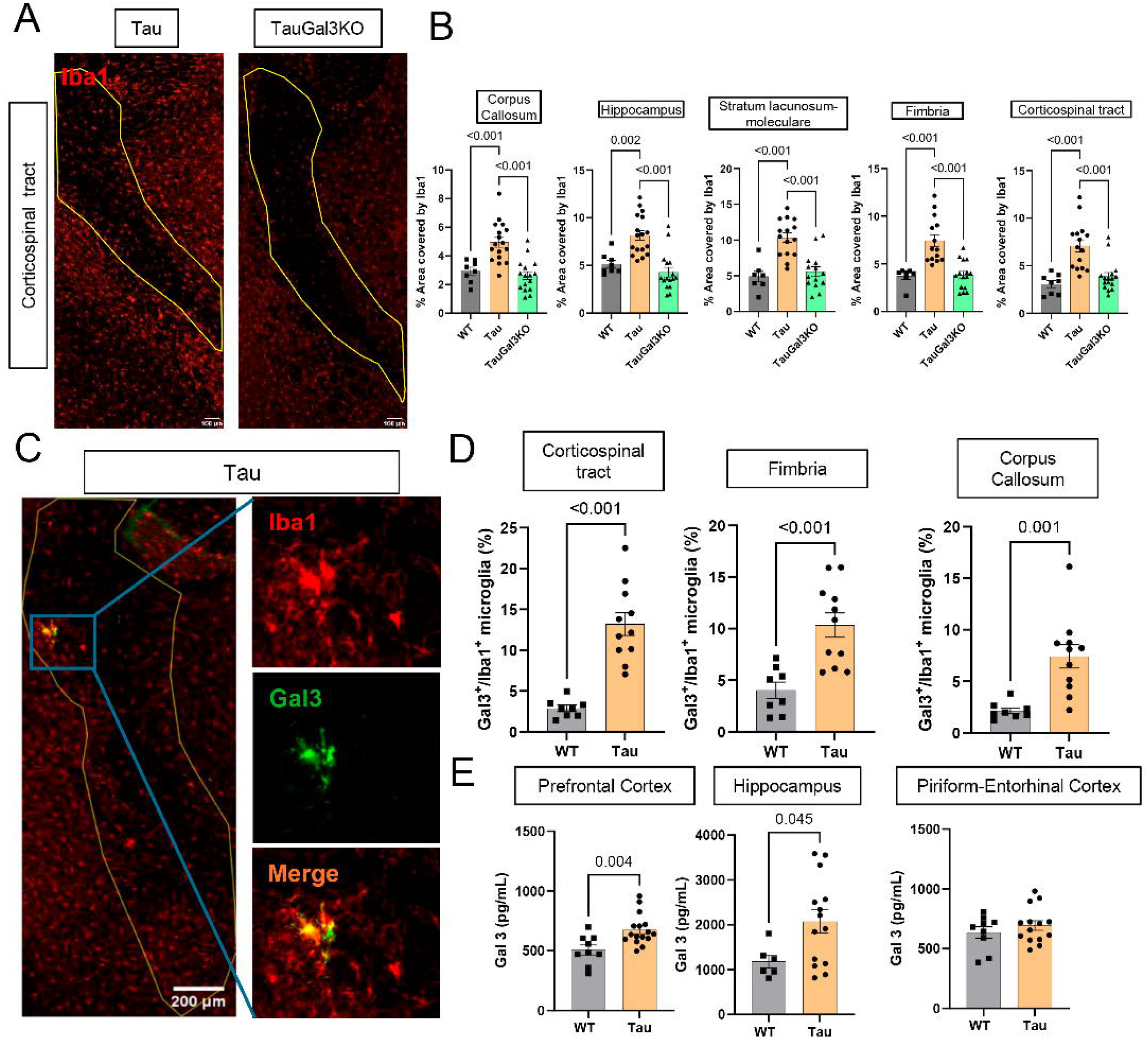
Galectin-3-dependent microglial activation and neuroinflammation in Tau mice. **(A)** Representative Iba1 immunostainings in Corticospinal tract region. **(B)** Quantification of percentage of area covered by Iba1 in gray and white matter areas (Corpus Callosum, Hippocampus, Stratum lacunosum-moleculare, Fimbria and Corticospinal tract). **(C)** Representative Iba1 and Gal3 immunostainings showing Gal3-positive (Gal3^+^) microglia in white matter in Tau mice. **(D)** Quantification of Gal3^+^ microglia normalized by total Iba1-positive (Iba1^+^) microglia in white matter areas. **(E)** ELISA quantification of Gal3 levels in RIPA-soluble gray matter areas (Prefrontal cortex, Hippocampus, Piriform-Entorhinal cortex). All values are expressed as individual experimental replicates with mean ±SEM. In B, one-ANOVA with Tukey’s multiple comparisons was performed. In D and E, unpaired t-test was performed. Significant p-values are shown.

### Galectin-3 deletion protects against tau-induced white matter degeneration and axonal pathology

White matter neurodegeneration is a key mechanism underlying the progression of frontotemporal dementia (FTD) (13, 38). In the Tau model, exacerbated neuroinflammation was associated with pronounced white matter degradation, a process that was markedly attenuated by Gal3 deletion. In the cortex, Tau mice displayed severe damage, manifesting as a drastic reduction in the percentage of myelinated cortex in Tau mice compared to WT (p<0.002. Fig. 3A). Contrary, the TauGal3KO mice were significantly protected from this damage, maintaining greater percentage of myelinated cortex (p=0.027) compared to the Tau group. Despite the reduction in the myelinated cortex, Tau mice displayed abnormal MBP coverage in white matter-enriched areas like the Corpus Callosum and the Stratum-Lacunosum-Moleculare (SLM) compared to WT mice (Fig. 3B-C). The latest was further confirmed by western blot, which demonstrates greater 20 kDa MBP levels in Tau mice compared to WT (Supp. Fig. 3A), indicating altered expression of this major exon-II-containing MBP isoform that is enriched in compact myelin and particularly sensitive to white matter injury (39). This anomalous MBP accumulation was prevented by Gal3 deletion in Tau, which displayed MBP levels similar to healthy control mice (Fig. 3B-C; Supp. Fig. 3A). Axonal integrity was also compromised in the Tau model, as indicated by a large percentage of axons showing vacuolar degeneration compared to WT in the Corpus Callosum and the Corticospinal tract (p<0.001, Fig. 3D). Consistent with the improved myelin integrity, TauGal3KO mice showed a significant reduction in vacuolar axonal degeneration compared to Tau mice (p<0.001; Fig. 3D). Furthermore, g-ratio analysis, which measures myelin thickness relative to axon diameter, confirmed that pathological changes observed in the Tau model were largely prevented in the TauGal3KO mice in the Corpus Callosum (p<0.001, Fig. 3F). Structural changes were also assessed using BlackGold Myelin staining. We found greater myelin density in Tau and TauGal3KO in the Corpus Callosum compared to control groups (Suppl. Fig. 3B), going along with the increased MBP levels detected (Fig. 3A; Suppl. Fig. 3A). Taking advantage of the BlackGold Staining, we also evaluate the axonal organization in the cortex. In this region, control mice exhibited lower coherency values, consistent with a more heterogeneous and multidirectional axonal organization (Suppl. Fig. 3C). In contrast, the pathological (Tau and TauGal3KO) group showed higher coherency, indicating a more uniformly aligned axonal pattern (Suppl. Fig. 3C). The increased coherency observed in the pathological mice may reflect a loss of structural complexity rather than improved organization.

**Fig. 3.**
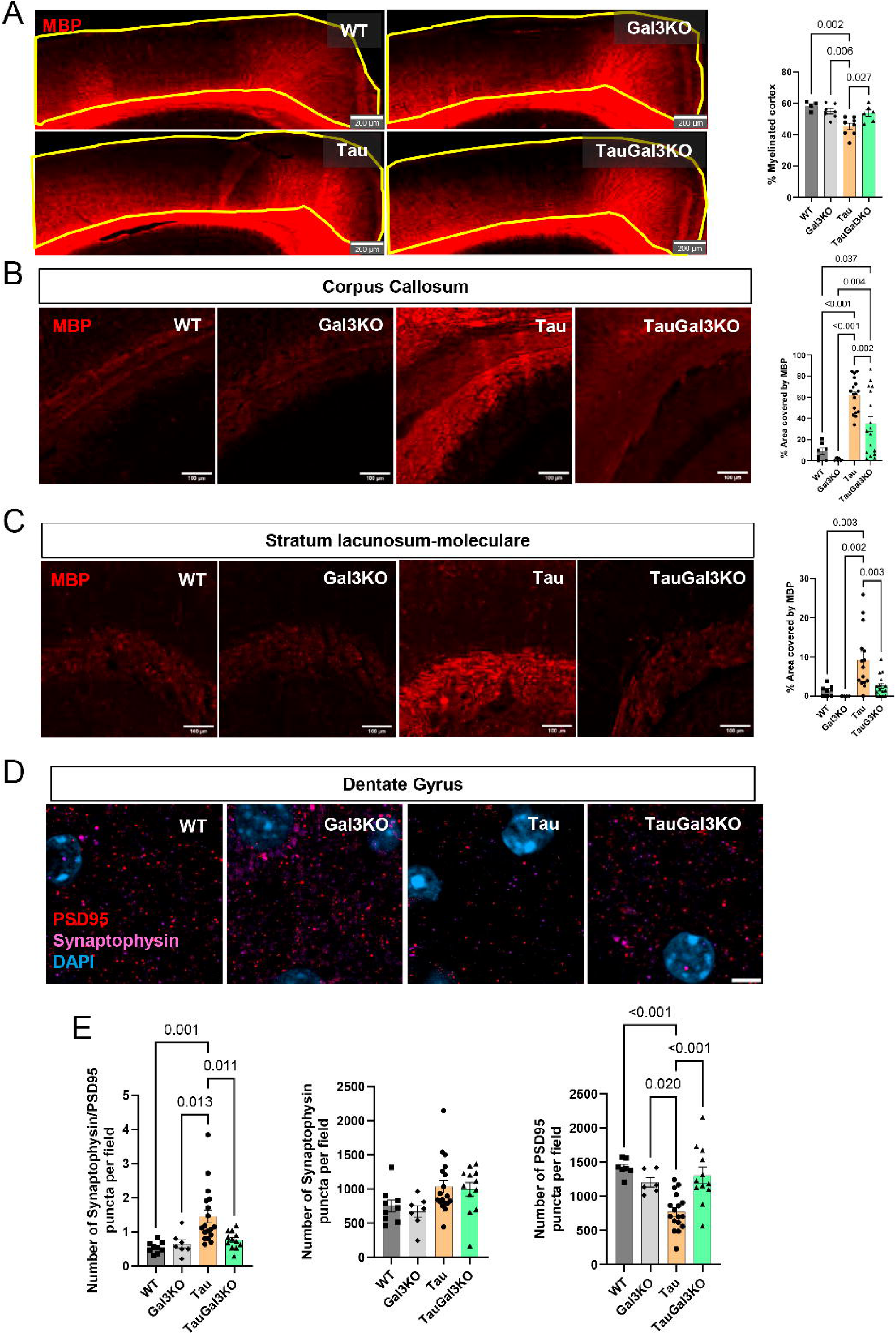
Galectin-3 deletion preserves white matter integrity and myelination in tauopathy. **(A)** Representative MBP immunostainings in Isocortex and quantification of the percentage of the cortex that presents myelination. **(B)** Representative MBP immunostaining in Corpus Callosum, with quantification of the percentage of area covered by MBP. **(C)** Representative MBP immunostaining in Stratum lacunosum-moleculare, with quantification of the percentage of area covered by MBP. **(D)** Representative transmission electron microscopy images of Corpus Callosum region. **(E)** Quantification of the percentage of axons presenting vacuolar degeneration in Corpus Callosum and Corticospinal tract regions. **(F)** Non-linear semilog correlation analysis of g-ratio and axon diameter across groups in Corpus Callosum and Corticospinal tract. In A, B, C and E, values are expressed as individual experimental replicates with mean ±SEM. In F, individual axon quantifications of Tau and TauGal3KO are shown in the graph, with non-linear fit curves and their 95% confidence interval of each group. In A, B, C and E, two-way ANOVA with Tukey’s multiple comparisons was performed. In F, extra sum-of-squares F-test was performed to calculate group effect of global comparison of fits, and pairwise difference between nonlinear regressions were assessed using extra sum-of-squares F-tests with Bonferroni correction for multiple comparisons. Significant p-values are shown.

### Tau pathology induces oligodendrocyte and white matter–associated microglial responses that are normalized by Galectin-3 deletion

In order to study if this myelin dysregulation was linked to oligodendrocytic levels and function, we studied these cells in white matter areas. The overall number of oligodendrocyte lineage cells (Olig2^+^) was significantly increased in Tau mice compared to WT/Gal3KO (p<0.001), potentially indicating a compensatory or reactive proliferation in response to the Tau-related myelin defects (Fig. 4A). The amount of mature myelinating oligodendrocytes, stained by CC1 (40), was also increased in Tau mice (Fig. 4B), which would support the greater MBP levels in white matter areas in Tau mice compared to WT. In Tau mice lacking Gal3, CC1 content in oligodendrocytes was lower, compared to WT mice, which is linked to the normalization of MBP levels in the Corpus Callosum and the SLM. In fimbria, CC1 content in oligodendrocytes showed a non-significant trend toward reduction in Tau mice lacking Gal3 compared with Tau mice (Suppl. Fig. 4A-B). These results were also accompanied by an increase in white matter microglia (WAM) signal in the Corpus Callosum of Tau mice, since the density Clec7a and Axl-positive microglia in these mice was higher than the rest of the groups (Fig. 4C).

**Fig. 4.**
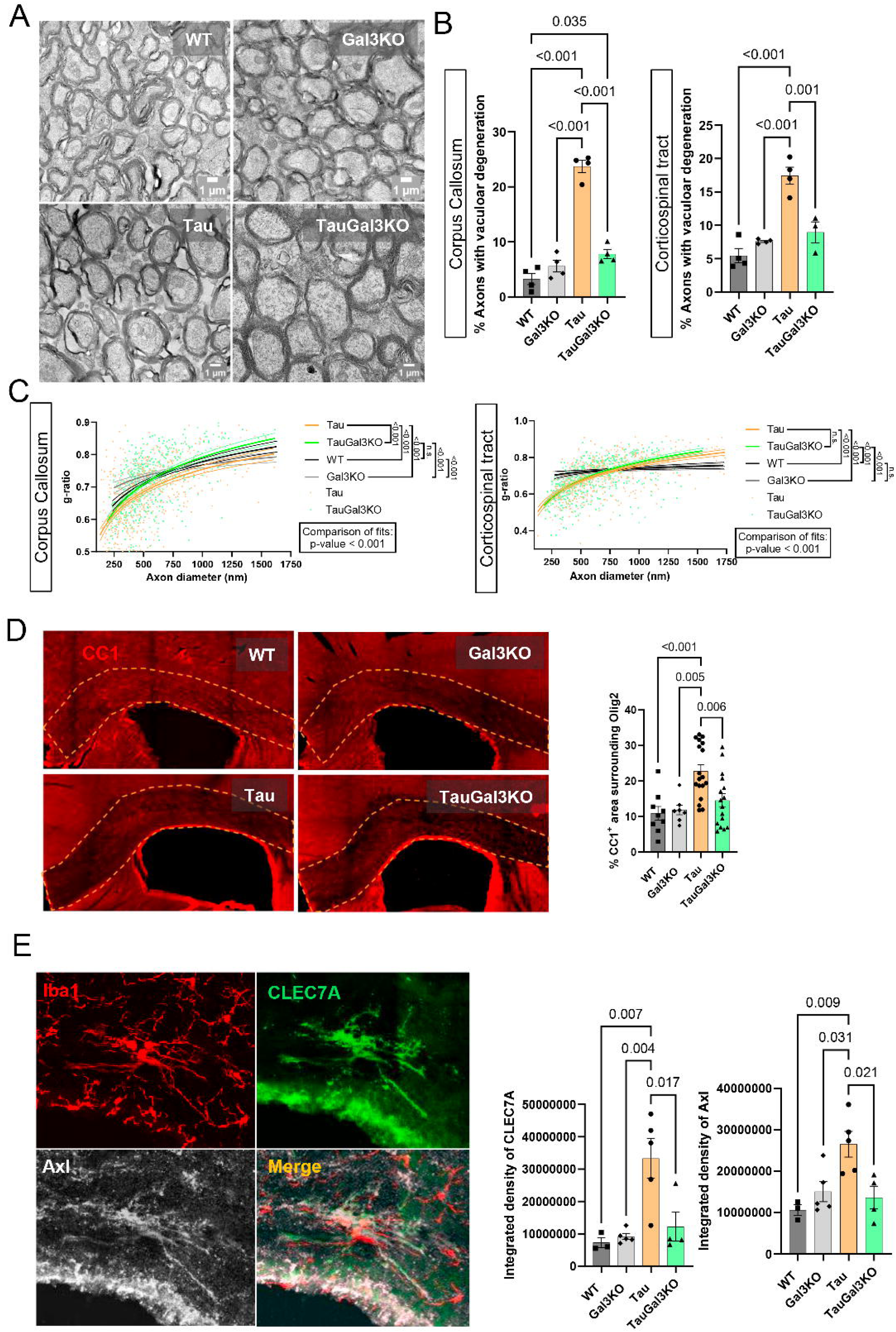
Galectin-3 deletion normalizes oligodendrocyte responses and white matter-associated microglial activation in tauopathy. **(A)** Representative Olig2 immunostainings in Corpus Callosum, with quantification of the number of Olig2^+^ oligodendrocytes per mm^2^. **(B)** Representative CC1 immunostainings in Corpus Callosum, with quantification of the CC1-positive (CC1^+^) area per Olig2^+^ oligodendrocytes. **(C)** Representative Iba1, Clec7a and Axl immunostaining in Corpus Callosum, with quantification of the integrated density of Clec7a and Axl. Values are expressed as individual experimental replicates with mean ±SEM. Two-way ANOVA with Tukey’s multiple comparisons was performed. Significant p-values are shown.

### Loss of Galectin-3 enhances microglial myelin phagocytosis and clearance

The protective effect observed upon Gal3 deletion could be linked to an enhanced capacity of microglia to interact and clear myelin debris, suggesting that Gal3 normally hinders efficient clearance mechanisms.

To investigate this, live-cell imaging was performed in BV2 microglial cells with (WT) or without galectin-3 (Gal3KO) following the addition of purified myelin. Over the 20-h imaging period, Gal3KO cells exhibited a higher proportion of myelin-positive cells and an increased amount of myelin uptake per cell compared with WT cells (Fig. 5A-B). In addition to enhanced uptake, myelin degradation was also increased in Gal3KO cells (Supplementary Fig. 5A). Furthermore, Gal3KO cells displayed increased lysosomal activation upon myelin exposure, as indicated by larger and more intensely fluorescent lysosomes (Supplementary Fig. 5B-D). To validate this result in the mouse model, the amount of MBP labelling within microglia in the Corpus Callosum was quantified. TauGal3KO mice showed a significantly increased percentage of Iba1 co-localizing with MBP compared to Tau mice (p=0.002; Fig. 5C). This increased co-localization indicates that, in the absence of Gal3, microglia exhibit enhanced phagocytic engagement with compromised myelin sheaths, reflecting an increased capacity for myelin clearance. The same *in vitro* system was utilized to study if this enhanced phagocytic microglial capacity displayed by Gal3KO BV2 cells was reflected in the context of tau pathology. Indeed, these cells displayed also higher fibrillar Tau uptake than their WT counterparts (Fig. 4D-E).

**Fig. 5.**
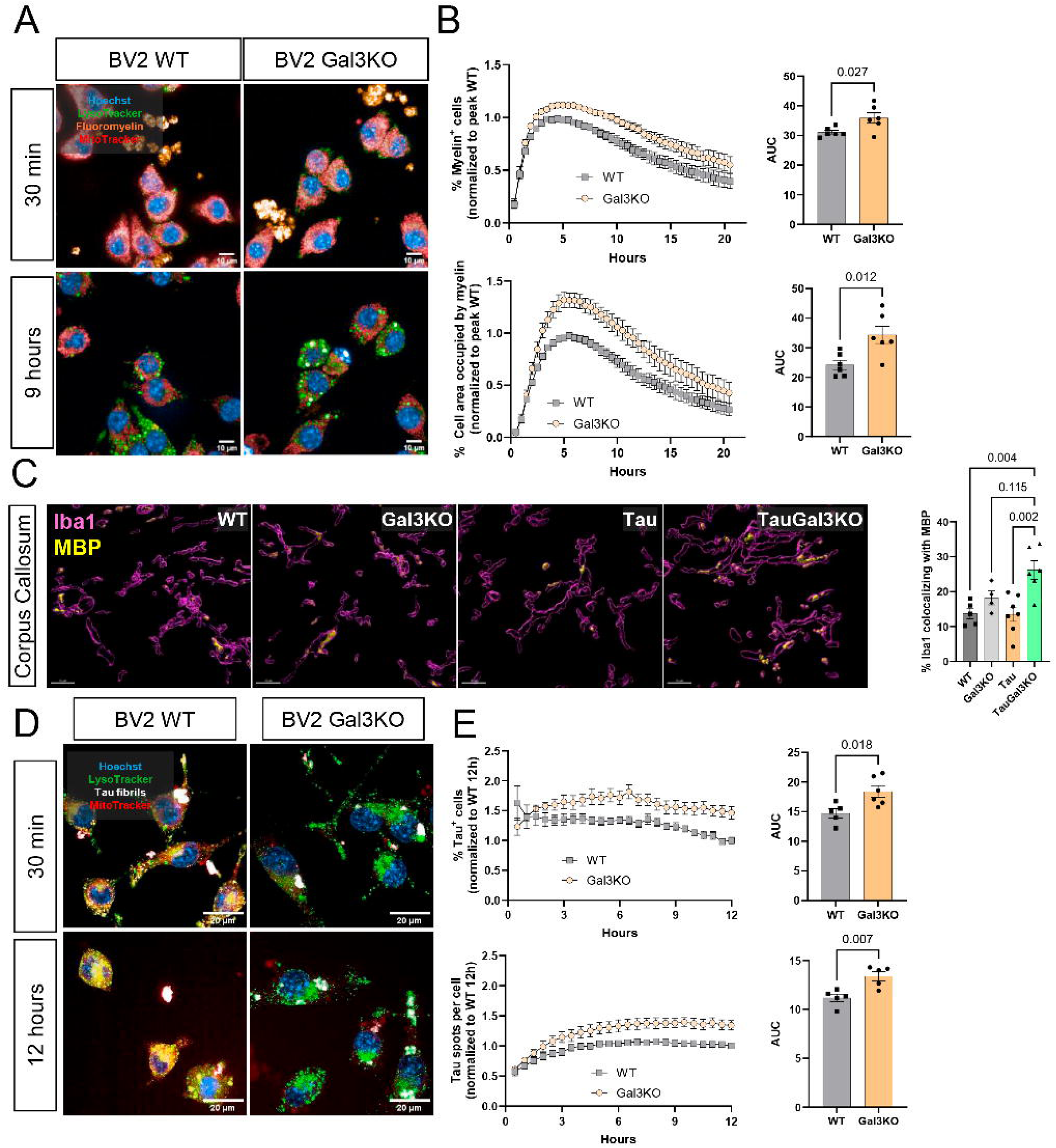
Galectin-3 deletion enhances microglial myelin and tau uptake. **(A)** Representative BV2 wild-type (WT) and Gal3 knockout (Gal3KO) images from live-cell imaging at 30 minutes and 9 hours after addition of myelin to cell cultures. **(B)** Quantification of live-cell imaging experiments showing percentage of cells with intracellular myelin (% Myelin^+^ cells, top) and amount of myelin per Myelin^+^ cell (% Cell area occupied by myelin, bottom) over time, with their area under the curve (AUC) quantifications. **(C)** Representative 3D reconstruction Imaris images of Iba1 and MBP immunostainings, with quantification of percentage of Iba1 colocalizing with MBP. **(D)** Representative WT and Gal3KO images from live-cell imaging at 30 minutes and 12 hours after addition of tau fibrils. **(E)** Quantification of live-cell imaging experiments showing percentage of cells with intracellular tau (% Tau^+^ cells, top) and amount of tau spots per cell (Tau spots per cell, bottom) over time, with their AUC quantifications. Values are expressed as individual experimental replicates with mean ±SEM. In B and E, unpaired t-test was performed. In C, two-way ANOVA with Tukey’s multiple comparisons was performed. Significant p-values are shown.

To assess how the interaction between tau and Gal3 affects brain cell populations, primary co-cultures of neurons, microglia, and astrocytes were established. Neuronal quantification revealed a Tau-Gal3-dependent toxicity, as the combined exposure to exogenous tau and Gal3 significantly reduced neuronal numbers. This effect was observed only when Gal3 was added after tau or when tau and Gal3 were preincubated and applied simultaneously, but not when either factor was applied alone. Notably, microglial numbers were significantly increased exclusively under these same conditions (Supplementary Fig. 5E-G).

### Galectin-3 deletion restores tau-associated proteomic alterations across hippocampal and entorhinal regions

Comparative pathway enrichment analysis revealed widespread proteomic dysregulation in Tau transgenic mice across multiple functional categories. In both the hippocampus (HC) and piriform-entorhinal cortex (EC), Tau mice exhibited a distinct proteomic profile characterized by extensive protein abundance changes relative to WT controls (Fig. 6A-B, Suppl. Fig. 6A). In contrast, the proteomic profile of TauGal3KO mice did not cluster with the Tau group, but instead aligned closely with WT and Gal3KO mice, indicating a broad normalization of Tau-associated proteomic alterations.

**Fig. 6.**
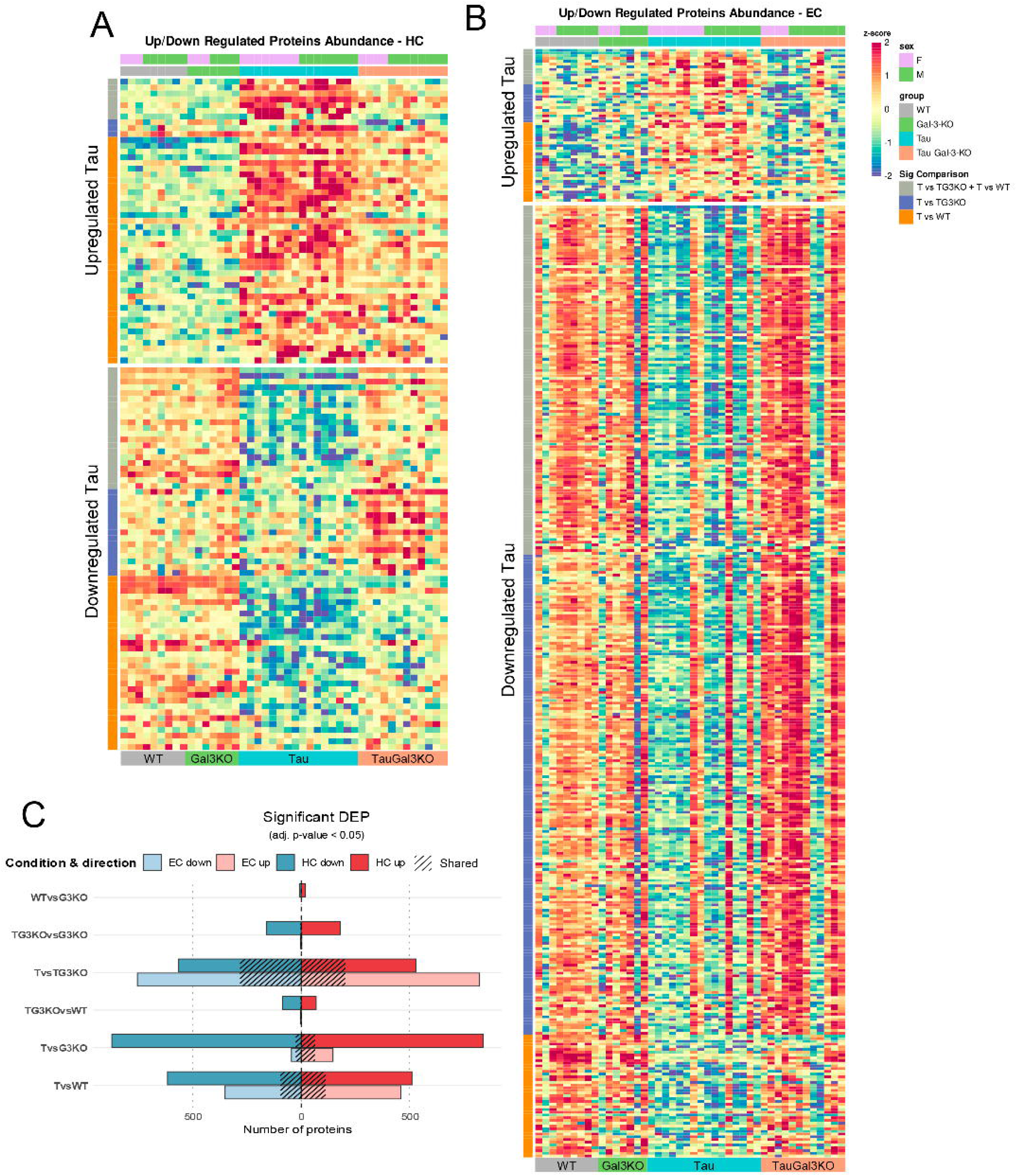
Galectin-3 deletion broadly normalizes tau-associated proteomic dysregulation in hippocampus and piriform-entorhinal cortex. **(A)** Heatmap of differentially-expressed proteins (DEP) in Hippocampus (HC) across groups, showing proteins upregulated and downregulated in Tau group. **(B)** Heatmap of differentially expressed proteins in Piriform-Entorhinal Cortex (EC) across groups, showing upregulated and downregulated in Tau group. **(C)** Quantification of the number of DEPs in HC and EC in the different comparisons, shared proteins within regions are marked with striped lines. T: Tau, TG3KO: TauGal3KO, G3KO:Gal3KO.

To quantify this effect, we analyzed the number of significantly differentially expressed proteins (DEPs) across experimental comparisons (Fig. 6C, Suppl. Table 1-2). The Tau vs. WT comparison revealed a large number of DEPs, reflecting extensive Tau-induced proteomic disruption. A similarly high number of DEPs was observed in the Tau vs. TauGal3KO comparison, indicating that TauGal3KO mice possess a proteomic profile distinct from Tau mice. Notably, direct comparison between TauGal3KO and WT mice revealed only a small number of DEPs. This reversal was further supported by Venn diagram analysis (Supplementary Fig. 6B), which demonstrated substantial overlap between proteins dysregulated in Tau vs. WT and those differentially expressed in Tau vs. TauGal3KO comparisons.

GSEA pathway enrichment analysis based on Reactome database identified adaptive immune response pathways as among the most significantly altered categories in both HC and EC (Fig. 7A-B). In Tau mice, components of the MHC class II antigen presentation pathway were strongly upregulated, and the differences were maintained when compared to TauGal3KO and WT.

**Fig. 7.**
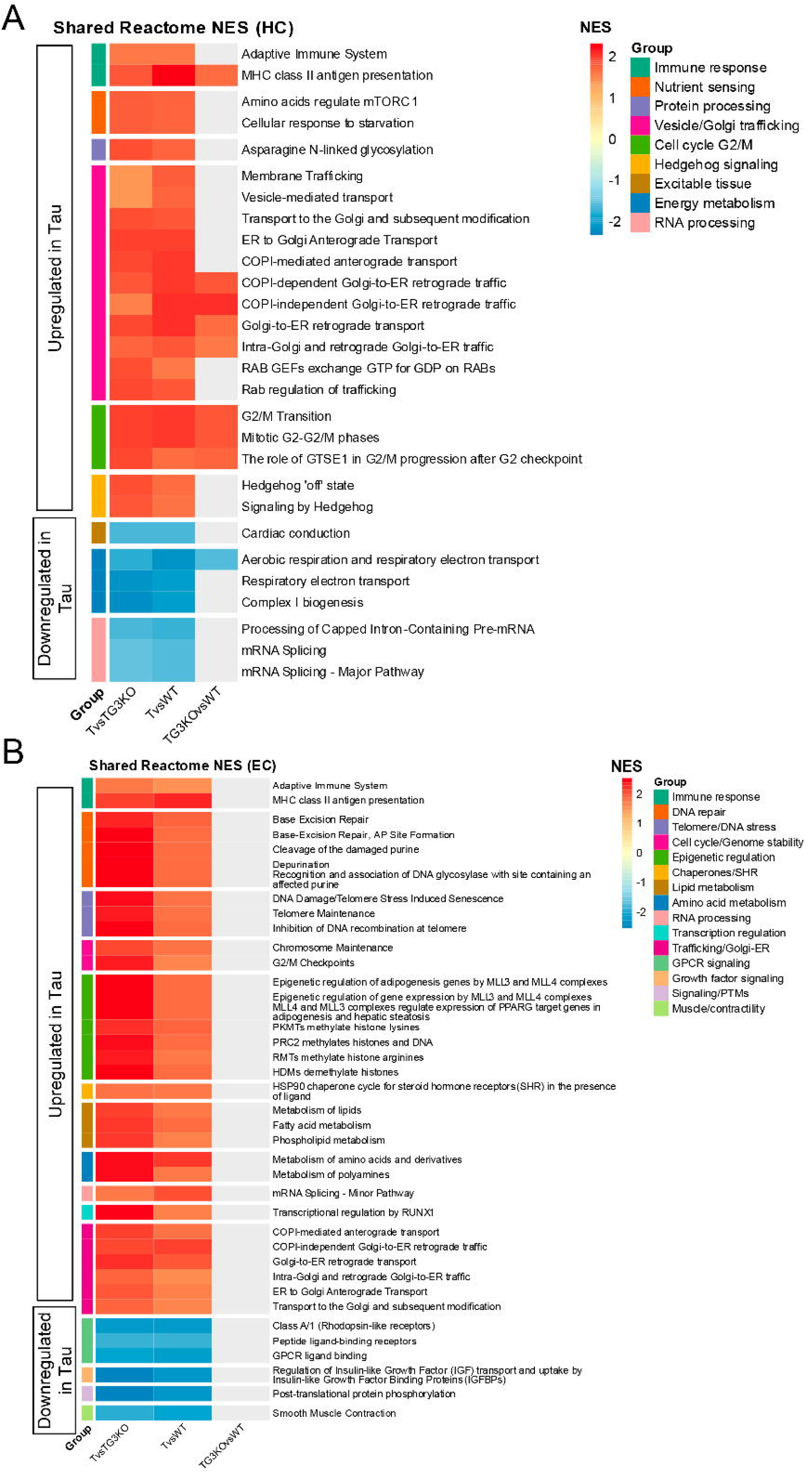
Galectin-3 deletion normalizes immune, vesicular trafficking, and mitochondrial pathway alterations in tauopathy. **(A)** Heatmap of changed Reactome pathways in Hippocampus (HC) across comparisons, showing pathways upregulated and downregulated in Tau group. **(B)** Heatmap of changed Reactome pathways in Entorhinal Cortex (EC) across comparisons, showing pathways upregulated and downregulated in Tau group. T:Tau, TG3KO: TauGal3KO, G3KO:Gal3KO.

Vesicular trafficking pathways were also prominently affected. In HC and EC, Tau mice showed significant upregulation compared to the other groups of ER-Golgi trafficking components, COPI- and Rab-associated proteins (Fig. 7A-B). Specifically in the EC, significant upregulation of lipid metabolism pathways was observed. In terms of the downregulated pathways restored, the most striking were mitochondrial metabolism pathways in the HC, such as respiratory electron transport.

KEGG database was utilized to validate these results, in which fatty acid degradation was upregulated and oxidative phosphorylation was downregulated in Tau, among others (Suppl. Fig. 6C, Suppl. Table 3). Curiously, KEGG enrichment identified a downregulation of SNARE interactions in vesicular transport, specifically in the EC of Tau mice compared to both TauGal3KO and WT, which likely reflects differential regulation of core fusion machinery versus upstream trafficking and contact-site scaffolds under tau-driven stress.

Phosphoproteomic analysis identified differential regulation in Tau vs TauGal3KO of multiple proteins associated with these pathways: Snx11, Rin1, Pacsin1, Q9ESE1, Ralgapa1, Gnao1, Ncor2, Mtpn and Xkr4 (Suppl. Fig. 7A). Moreover, proteomic quantification of tau-related kinases identified several enzymes implicated in Tau-Gal3 interactions, including Mapk8, Mapk10, Csnk1d, Csnk1e, Fyn, Akt1, and Akt2 (Supplementary Fig. 7B). Phosphorylation site analysis of all tau-related kinases revealed several targets in Tau mice compared with WT, Gal3KO, and TauGal3KO groups, with the most prominent changes observed in the hippocampus. Differentially regulated phosphorylation sites included Gsk3β S389 and S9, Akt3 T447, Braf S735, Prkaa1 S356, Brsk1 T477 and S311, and Prkacb S339 (Suppl. Fig. 7C).

Given that one of the most prominently altered proteins was the GABA receptor subunit Gabra2, together with previous evidence implicating Gal3 in interneuronal dysfunction during neurodegeneration (41) and the known vulnerability of interneurons, particularly parvalbumin-positive cells, to impairments in oxidative phosphorylation (42), we hypothesized that the observed proteomic alterations extend beyond excitatory neurons to also affect inhibitory interneurons. We therefore quantified the amount of mitochondrial complex I subunit NDUFB8 in MAP2-positive (MAP2^+^) or parvalbumin-positive (PV^+^) cells in hippocampus. Indeed, TauGal3KO mice showed a significant increase in NDUFB8 in both MAP2^+^ and PV^+^ cells (Suppl. Fig. 8A), which is linked to proper mitochondrial function (43, 44).

In order to bring this murine-related effects to a human context, IPS-derived neurons, either wild-type (WT) or with P301L mutation (Tau) were studied (Suppl. Fig. 8C). In this context, we first studied if Gal3 addition to these human cells impacted tau pathology post-seeding with tau fibrils. Indeed, giving Gal3 after seeding with tau fibrils increased the amount of AT8-positive (AT8^+^) spots per neuron (Suppl. Fig. 8C, Experiment 1). We then quantified if this Gal3-provoked tau hyperphosphorylation could be prevented by addition of Gal3 inhibitor TD139, and indeed this inhibitor decreased the amount of AT8+ spots per neuron (Suppl. Fig. 8C, Experiment 2-3). Finally, we characterized the extent of hyperphosphorylated tau to be also aggregated, by utilizing the compound HS84. We identified a decrease in aggregated (HS84-positive) and hyperphosphorylated (PHF1^+^) tau after the addition of TD139 inhibitor (Suppl. Fig. 8D, Experiment 4). Not only that, but we also observed a change in NDUFB8 content per neuron, after Gal3 inhibitor was added.

Together, these findings indicate that Gal3 critically shapes tau-driven proteomic, signaling, and metabolic alterations in both murine and human neuronal systems, with effects extending from vesicular trafficking and immune pathways to mitochondrial function in excitatory and inhibitory neuronal populations; establishing Gal3 as a central modulator of tau-driven molecular and cellular dysfunction.

## Discussion

Frontotemporal dementia (FTD) represents a clinically and pathologically heterogeneous group of neurodegenerative disorders characterized by the progressive atrophy of the frontal and temporal lobes. Among the genetic causes of FTD, mutations in the *MAPT* gene are a primary driver of disease, leading to the abnormal aggregation of tau protein—a condition collectively termed tauopathy. Pathogenic *MAPT* mutations promotes tau species accumulate as neurofibrillary tangles and glial inclusions, which correlate strongly with synaptic loss, neuronal death, white matter alterations and the distinct patterns of cortical atrophy seen in patients (45, 46).

Emerging evidence suggests that neuroinflammation is a central driver that may even precede overt protein aggregation (5). In the context of *MAPT*-dependent tauopathy, microglia transition from a homeostatic state to a reactive phenotype, creating a feedback loop where inflammatory signaling exacerbates tau hyperphosphorylation and spread, while accumulating protein aggregates further stimulate microglial reactivity. Understanding the molecular triggers that orchestrate this neuroinflammatory environment is essential for developing therapeutic strategies. The findings presented in this study establish Gal3 as a central upstream regulator that orchestrates the neuroinflammatory environment driving both Tau pathogenesis and associated white matter neurodegeneration.

By utilizing a Gal3-deficient tauopathy model, we demonstrate that its absence leads to a systemic attenuation of core pathological markers. Our histological analysis reveals that Gal3 deletion significantly reduces the load of hyperphosphorylated Tau (AT8^+^) and pathological Tau conformations (MC1) in regions highly vulnerable to disease progression, including the isocortex, piriform-entorhinal cortex, and hippocampus (46). Interestingly, the reduction in cortical AT8 load was particularly prominent in female mice lacking Gal3, suggesting a potential sex-specific vulnerability to Gal3 modulation that warrants further investigation. These histological findings are strongly corroborated by biochemical evidence from hippocampal homogenates, which showed significant decreases in toxic Tau species, including early phosphorylation markers such as pTau231, as well as established pathological species like pTau217 and pTau181. Increased kinases protein abundance and differential regulation across multiple phosphorylation sites underscores the role of Gal3 in fostering a pro-aggregatory environment, likely triggering microglial interaction through its known relation to immune receptors like TLR4 and TREM2 (24, 25).

Beyond proteinopathy, white matter alterations and neuroinflammation, which are recognized as a key drivers of clinical decline in Frontotemporal Dementia (FTD) (15, 38, 47). Clinical studies highlighted the importance of inflammatory response and microglial activation in FTD patients (10, 48, 49) in the progression of FTD and Diffusion Tensor Imaging has demonstrated significant white matter microstructural integrity alterations among FTD patients, particularly in regions such as the corpus callosum (15). Furthermore, Tau deposition in cortical areas has been shown to induce secondary atrophy in connected white matter regions (50). The exacerbated neuroinflammation associated with tauopathy was characterized in our model by high Gal3 expression in activated microglia within white matter tracts, such as corticospinal tract or corpus callosum. Microglial activation was also characterized by the expression of Clec7a and Axl, both microglial markers linked to white matter-associated microglia (WAM) (19), in the corpus callosum. The percentage of myelinated cortex was profoundly lower in Tau mice relative to wild-type animals, a hallmark of the extensive white matter degeneration driven by tauopathy (51). Along with that, Tau mice suffered from severe vacuolar axonal degeneration and pathological myelin sheets surroundings axons (g-ratio) in the CC. Beyond this axonal loss, there was also an abnormal accumulation of CC1+ mature myelinating oligodendrocytes in the corpus callosum and a corresponding anomalous increase in MBP coverage, specifically the 20 kDa MBP isoform.

Crucially, the genetic deletion of Gal3 largely prevented these structural defects, maintaining axonal integrity and g-ratios comparable to healthy controls in the corpus callosum and the corticospinal tract. The absence of Gal3 normalized both the number of mature oligodendrocytes and the biochemical levels of the 20 kDa MBP isoform in the corpus callosum. These findings align with the recently established role of microglial cells as essential guardians for the maintenance of myelin sheaths and the prevention of age- or pathology-related degeneration (20). Our results suggest that by removing the Gal3, the homeostatic capacity of microglia to clear myelin debris is restored, providing a robust neuroprotective effect. This neuroprotective effect is also sustained by the maintenance of the homeostatic oligodendrocyte activity, which prevents the unproductive accumulation of mature cells, thereby ensuring functional axonal myelination and safeguarding axonal integrity

A pivotal mechanism identified in this study is the enhanced capacity of microglia to clear tau aggregates and myelin debris in the absence of Gal3. Both *in vitro* live-cell imaging and *in vivo* colocalization data demonstrate that Gal3 deficiency leads to increased tau and myelin uptake and degradation myelin debris. In the corpus callosum, the significantly increased colocalization of Iba1 with MBP in TauGal3KO mice indicates a more effective phagocytic engagement with compromised myelin sheaths. This evidence supports the concept that Gal3 may normally act as a “molecular brake” that hinders homeostatic clearance mechanisms. By removing Gal3, microglia appear to regain their ability to clear the niche, thereby preventing the secondary axonal damage and lysosomal dysfunction typically observed under tau-driven stress.

The systemic impact of Gal3 deletion is further evidenced by proteomic profiling, which revealed a broad normalization of Tau-induced alterations across the hippocampus and entorhinal cortex. Notably, the extensive proteomic disruption of vesicular trafficking and lipid metabolism pathways seen in Tau mice was effectively normalized in the TauGal3KO model. Given that lipid alterations are a recognized cause of oligodendrocyte malfunction (52), this proteomic stabilization underscores the role of Gal3 in maintaining the metabolic environment necessary for white matter preservation. Most notably, Gal3 deletion also restored oxidative phosphorylation pathways and mitochondrial health. Specifically, TauGal3KO mice displayed a significant increase in the mitochondrial complex I subunit NDUFB8 in both excitatory neurons and parvalbumin-positive (PV+) interneurons. Given the known metabolic vulnerability of PV+ cells to oxidative stress (43, 44), this restoration suggests that Gal3 inhibition may protect the brain’s essential inhibitory circuitry from tau-driven mitochondrial alterations.

Finally, the translational relevance of these findings was validated in human iPSC-derived neurons, where exogenous Gal3 exacerbated Tau hyperphosphorylation following fibril seeding, an effect that was effectively reversed by the Gal3 inhibitor TD139. Furthermore, TD139 treatment reduced the accumulation of aggregated Tau and normalized mitochondrial markers in these human cells. Collectively, these results establish Gal3 as a central modulator of tau-driven molecular and cellular dysfunction. By targeting Gal3, it may be possible to simultaneously dampen detrimental neuroinflammation, enhance the clearance of toxic protein aggregates, and protect the metabolic integrity of the brain, making it a highly promising therapeutic target for halting the progression of frontotemporal dementia and related tauopathies.

## Acknowledgments

We thank Sara Linse and Emil Axell, Lund University, for the purification of tau proteins. Thanks to the Cell and Gene Therapy Core (Lund Stem Cell Centre) for the design of the CRISPR-mediated KO experiment. We thank the Lund Protein Production Platform (LP3, www.lu.se/lp3), Lund University, for the production of the recombinant human Gal3. Lund University Bioimaging Centre (LBIC), Lund University, is gratefully acknowledged for providing experimental resources.

## Authors’ contributions

Conceptualization, L.C.-F., J.F.-R., A.B.-S., J.L.V. and T.D.; methodology, L.C.-F., J.S.-H., A.L.-A., L.R.R., K.P., E. A., E.V., Y.Y.; investigation, L.C.-F., M.D’E., J.S.-H., A.L.-A., L.R.R., J.F.-R., K.P.; writing original draft, L.C.-F and A.B.-S.; funding acquisition, T.D., J.L.V., A.B.-S., and J.V.; supervision, J.G.-R., J.L.V., A.B.-S., J.F.-R. and T.D. All authors reviewed the manuscript.

