## Supporting information for "Galectin-3 drives tau-associated neuroinflammation, white matter degeneration and proteomic dysregulation"

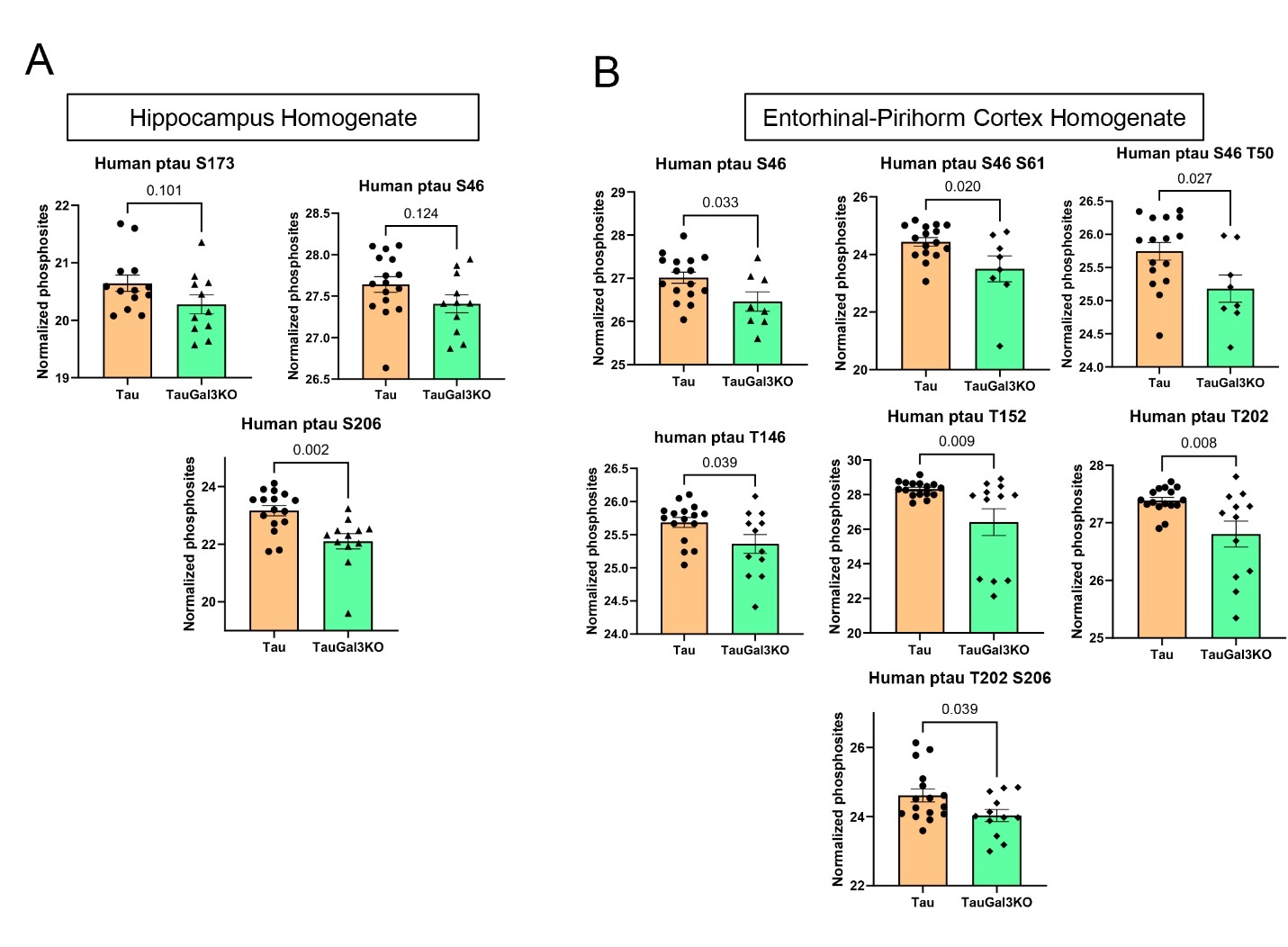

**Suppl. Fig. 1. Galectin-3 deletion alters tau phosphorylation profiles in RIPA-soluble hippocampal and entorhinal-piriform cortex fractions.**

**(A)** Phosphoproteomic quantification of human tau-related phosphorylations changed between Tau and TauGal3KO in RIPA-soluble hippocampi homogenates. **(B)** Phosphoproteomic quantification of human tau-related phosphorylations changed between Tau and TauGal3KO in RIPA-soluble entorhinal-piriform cortex homogenates. All values are expressed as individual experimental replicates with mean ±SEM. Unpaired t-test was performed. Significant p-values are shown.

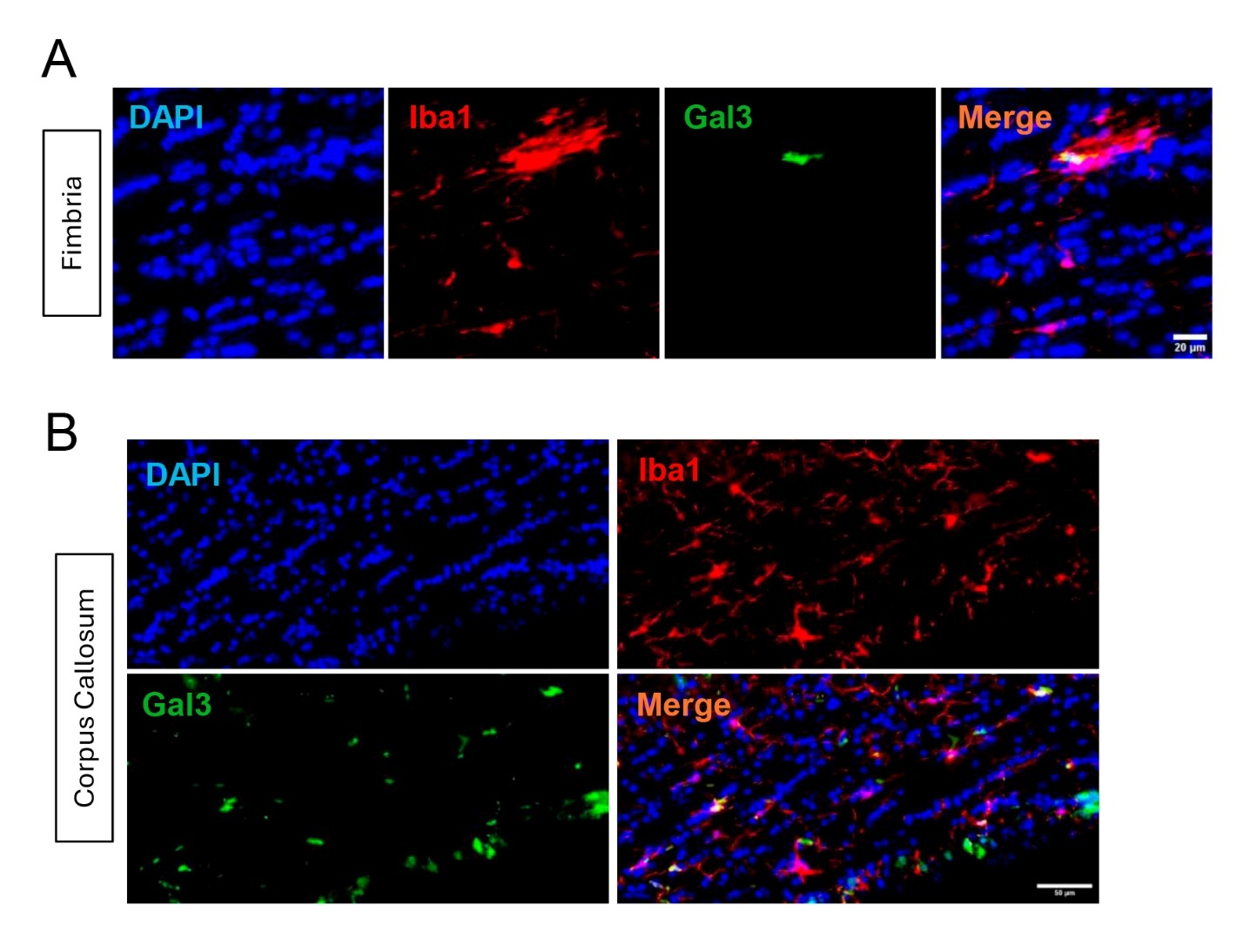

**Suppl. Fig. 2. Galectin-3-positive microglia are enriched in white matter tracts of Tau mice.**

**(A)** Representative Iba1 and Gal3 immunostainings showing Gal3-positive (Gal3^+^) microglia in Fimbria of Tau mice. **(B)** Representative Iba1 and Gal3 immunostainings showing Gal3^+^ microglia in Corpus Callosum of Tau mice.

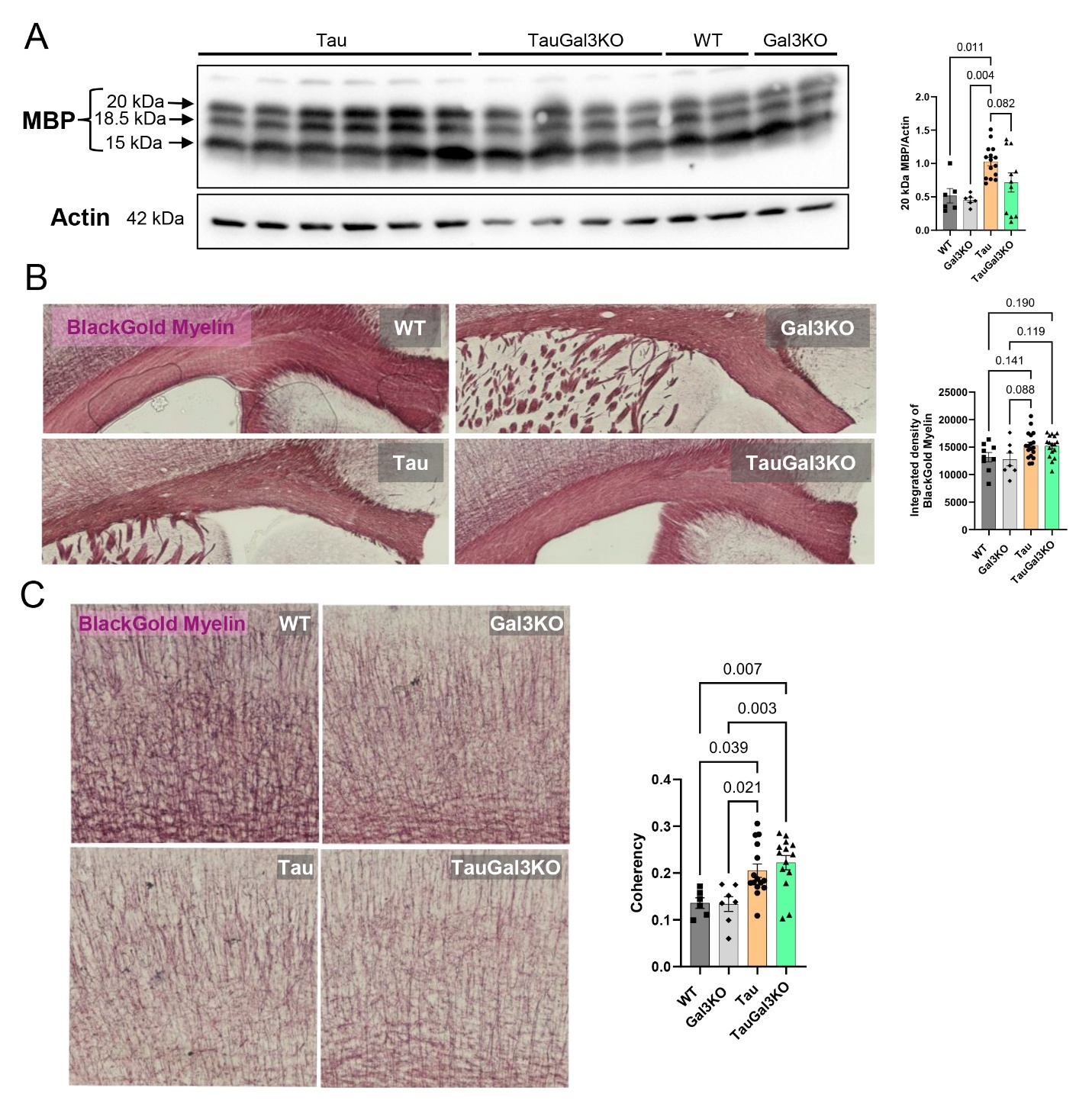

**Suppl. Fig. 3. Altered myelin basic protein levels and myelin organization in tauopathy.**

(A) Western blot quantification of myelin basic protein (MBP) related to Actin in RIPA-soluble hippocampi homogenates. (B) Representative BlackGold Myelin staining of Corpus Callosum and quantification of integrated density of the dye. (C) Representative BlackGold Myelin staining of cortical fibers and quantification of their coherency values. All values are expressed as individual experimental replicates with mean ±SEM. Two-way ANOVA with Tukey’s multiple comparisons was performed. Significant p-values are shown.

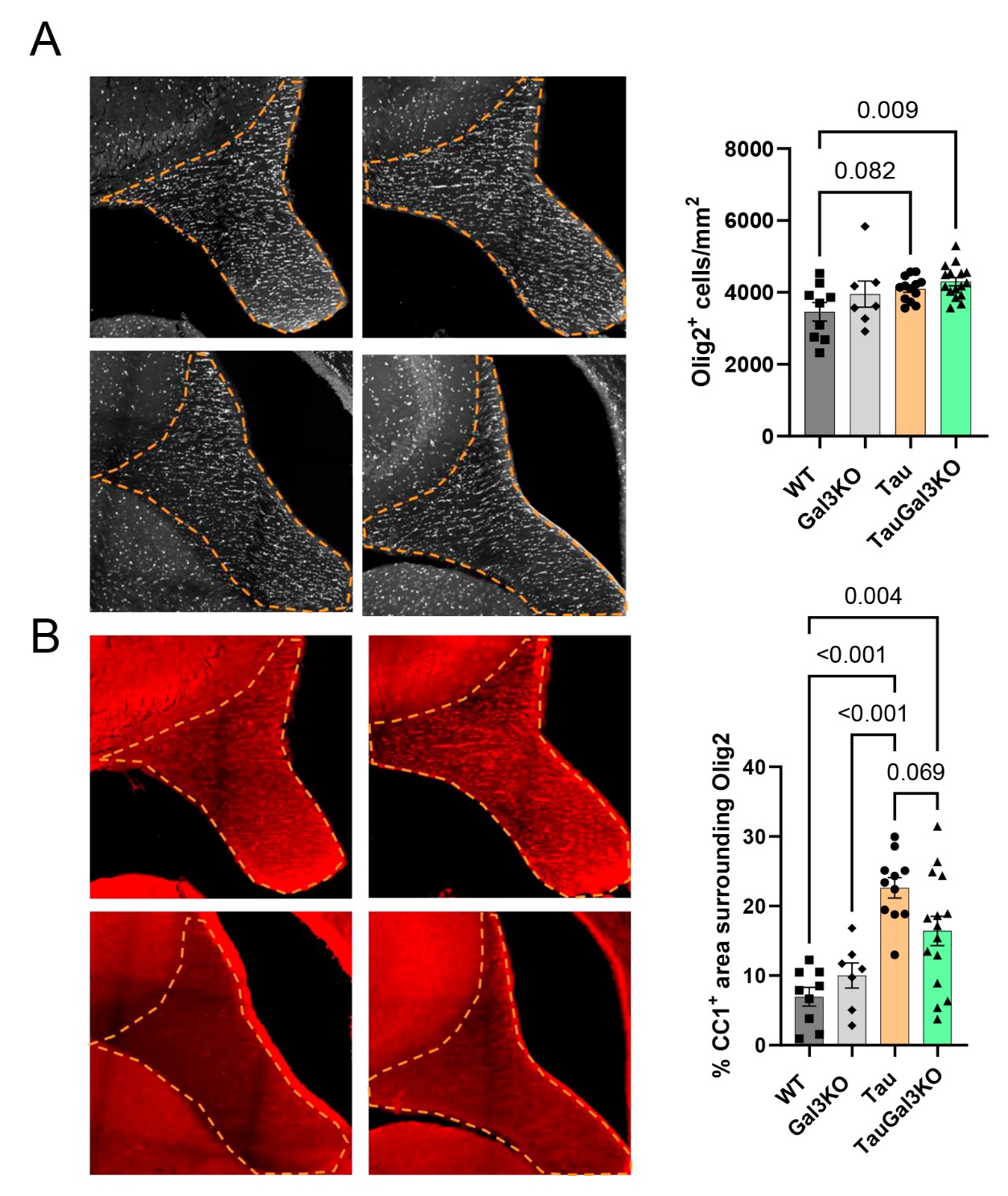

**Suppl. Fig. 4. Oligodendrocyte lineage and maturation markers in the fimbria of tauopathy mice.**

**(A)** Representative Olig2 immunostainings in Fimbria, with quantification of the number of Olig2^+^ oligodendrocytes per mm^2^. **(B)** Representative CC1 immunostainings in Fimbria, with quantification of the CC1-positive (CC1^+^) area per Olig2^+^ oligodendrocytes. Values are expressed as individual experimental replicates with mean ±SEM. Two-way ANOVA with Tukey’s multiple comparisons was performed. Significant p-values are shown.

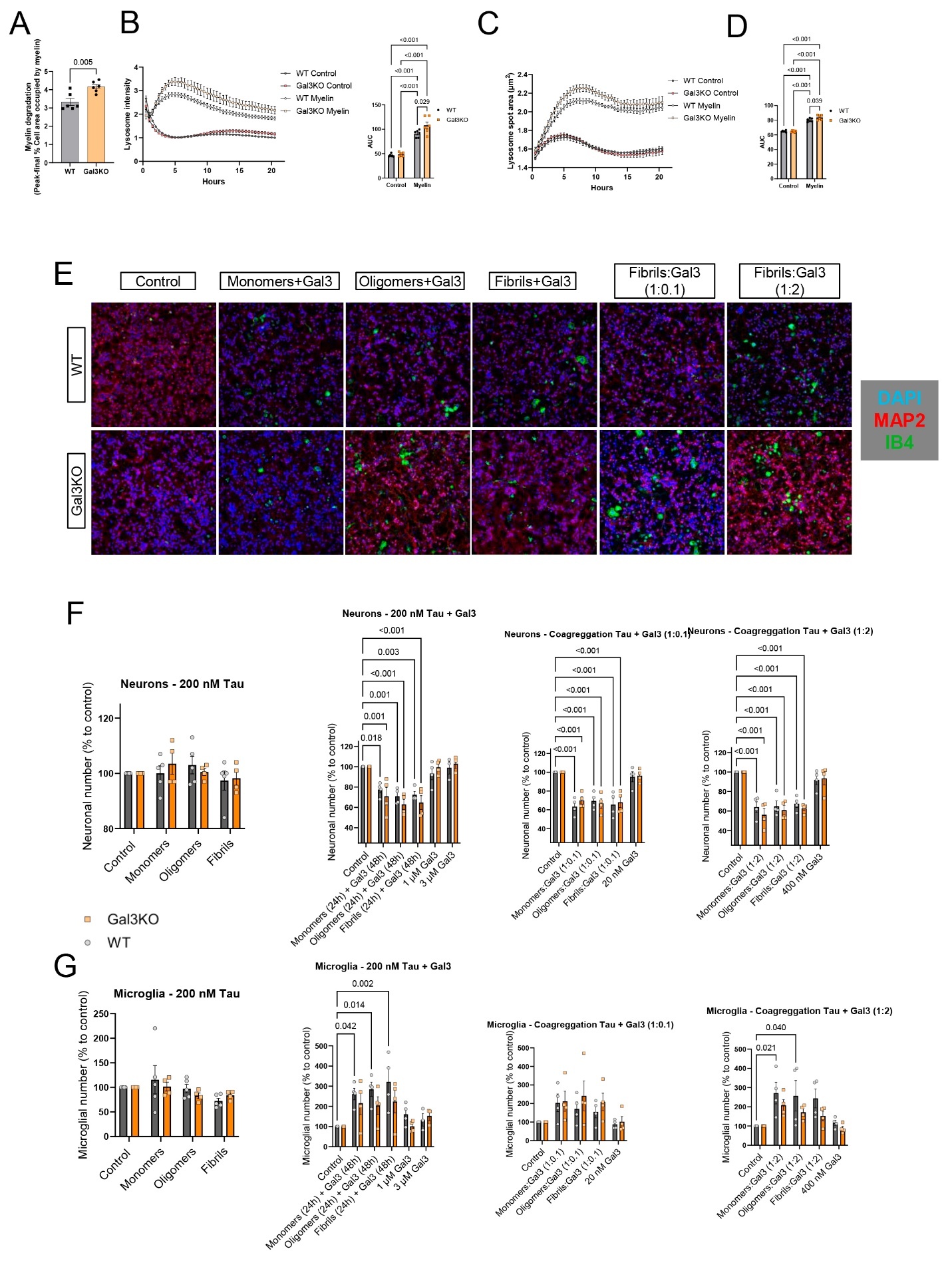

**Suppl. Fig. 5. Enhanced myelin degradation, lysosomal activity, and cell-type–specific responses following Galectin-3 modulation**

**(A)** Quantification of BV2 myelin degradation from live-cell imaging experiments, measured by the maximum amount of intracellular myelin per condition (Peak), minus the final measurement. **(B)** Quantification of LysoTracker particle intensity in live-cell imaging experiments with area under the curve (AUC) quantification over time. **(C,D)** Quantification of LysoTracker particle area with AUC quantification over time. **(E)** Representative immunostaining of primary cocultures with MAP2 and isolectin-B4 (IB4). **(F)** Quantification of the number of neurons throughout the different coculture experimental conditions, normalized to control numbers. **(G)** Quantification of the number of microglia throughout the different coculture experimental conditions, normalized to control numbers. Values are expressed as individual experimental replicates with mean ±SEM. In A, unpaired t-test was performed. In B, D, F and G, two-way ANOVA with Tukey’s multiple comparisons was performed. Significant p-values are shown.

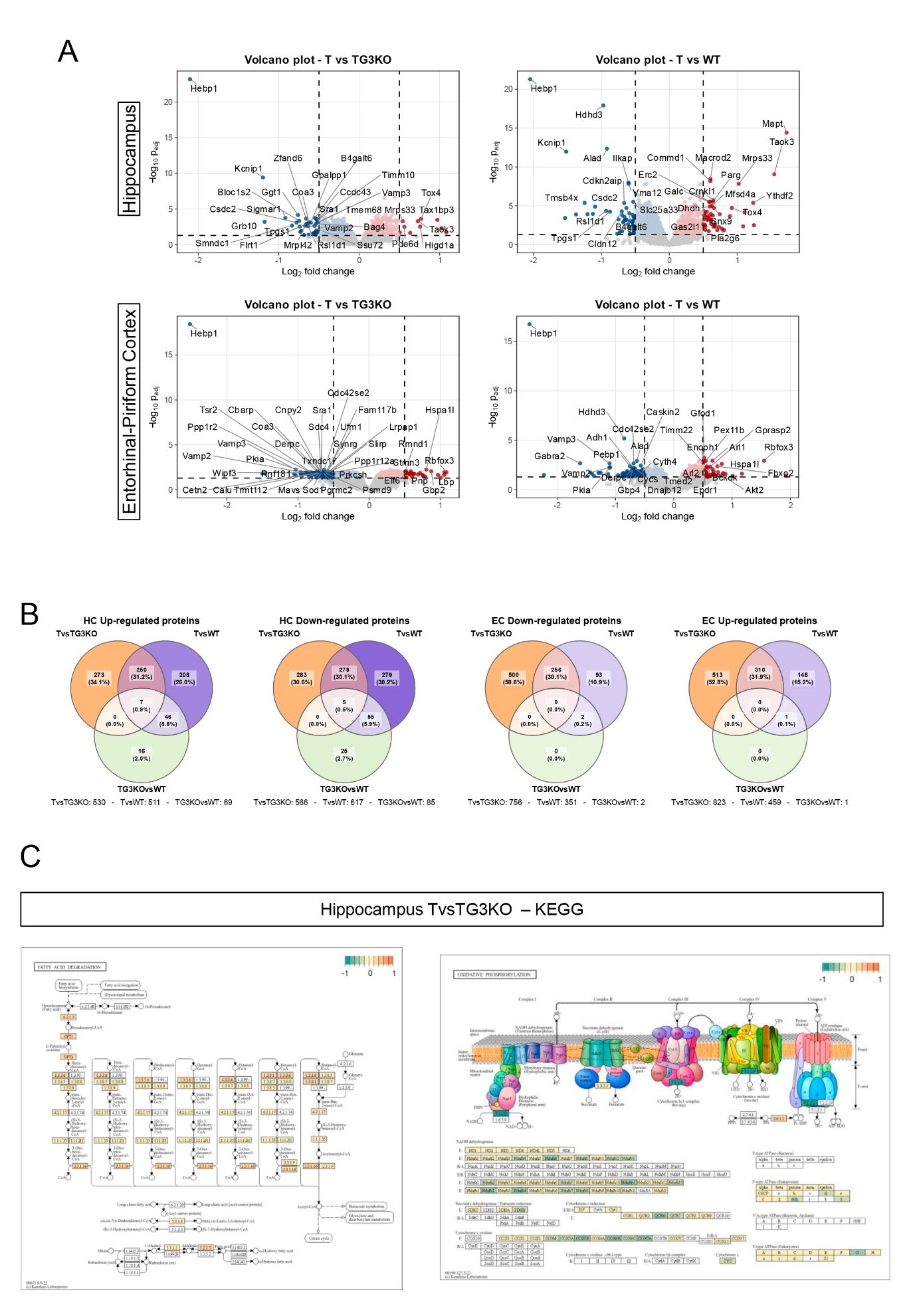

**Suppl. Fig. 6. Galectin-3 deletion reverses tau-associated proteomic alterations in hippocampus and entorhinal cortex**

**(A)** Volcano plot showing the most differentially expressed proteins in Tau versus TauGal3KO and Tau versus WT comparisons, in hippocampus (HC) and piriform-entorhinal cortex (EC). **(B)** Venn diagrams showing the shared proteins between the different comparisons. **(C)** KEGG diagrams showing the protein changes in Tau versus TauGal3KO in fatty acid degradation and oxidative phosphorylation pathways in hippocampus. T: Tau, TG3KO: TauGal3KO, G3KO:Gal3KO.

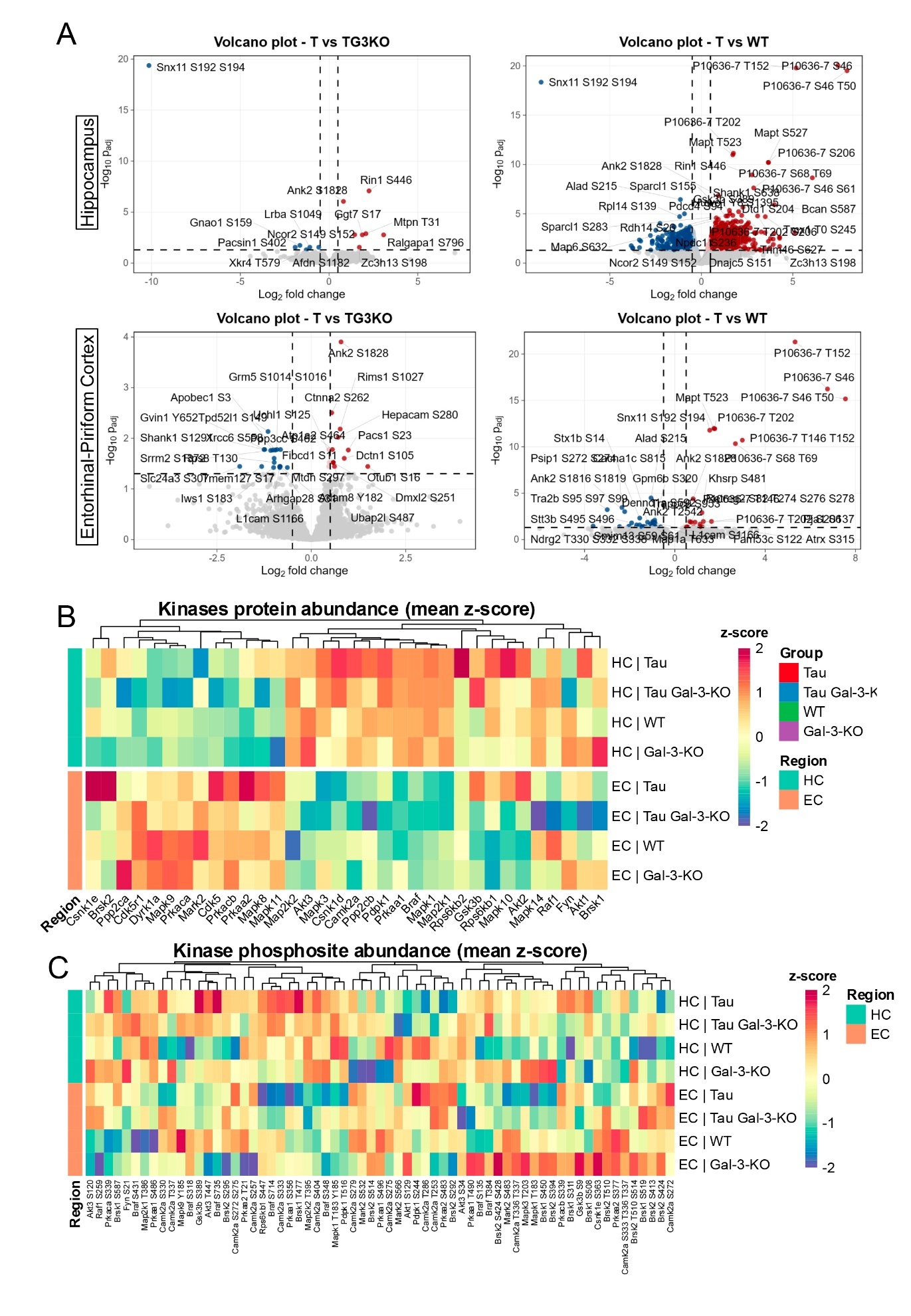

**Suppl. Fig. 7. Galectin-3 deletion modulates tau-associated kinase abundance and phosphorylation states**

**(A)** Volcano plot showing the most differentially expressed phosphosites in Tau versus TauGal3KO and Tau versus WT comparisons, in hippocampus (HC) and piriform-entorhinal cortex (EC). **(B)** Heatmap showing the protein abundance of tau-related kinases. **(C)** Heatmap showing the phosphosites abundance of the tau-related kinases. T: Tau, TG3KO: TauGal3KO, G3KO:Gal3KO.

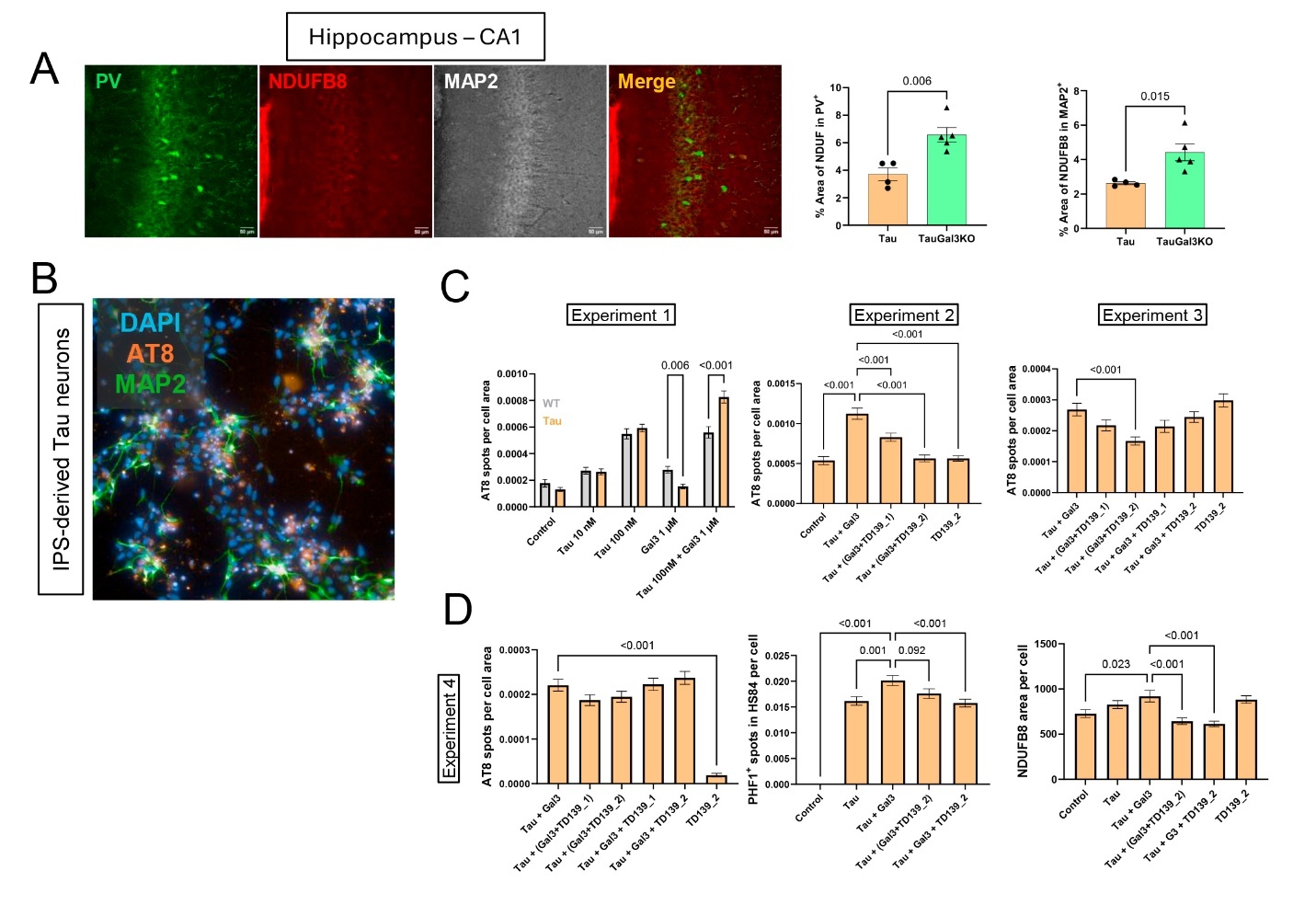

**Suppl. Fig. 8.** **Galectin-3 modulation affects neuronal mitochondrial integrity and tau pathology in mouse hippocampus and human iPSC-derived neurons.**

**(A)** Representative parvalbumin (PV), NDUFB8 and MAP2 immunostainings in CA1 region of HC, with quantification of NDUFB8 area within PV and MAP2 cells. **(B)** Representative immunostaining of IPS-derived P301L mutated neurons (Tau neurons) showing AT8 and MAP2. **(C)** Experiments 1-3 showing individual neuronal quantification of the amount of AT8-positive (AT8^+^) spots per cell throughout the experimental conditions. **(D)** Experiment 4 with AT8^+^ intraneuronal quantification, as well as amount of aggregated (HS84) and phosphorylated (PHF1) sites and mitochondrial complex I (NDUFB8) content per neuron. In A, values are expressed as individual experimental replicates with mean ±SEM. In C and D, values are expressed as mean ±SEM, and individual replicates signifying one neuron (>300 neurons per conditions were quantified). In A, unpaired t-test was performed. In C and D, one-way ANOVA was performed with Tukey’s multiple comparisons. Significant p-values are shown.

**Suppl. Table S1. Differential protein expression and pathway enrichment analyses of hippocampal proteomic data**. This table comprises five sheets (A–E). **A)** contains the results of differential expression analysis performed using the *limma* framework on hippocampal proteomic data, including proteins that passed quality control and filtering criteria. For each protein, log2 fold change (logFC), raw *p* value, and multiple-testing adjusted *p* value (adjusted.p.value; Benjamini–Hochberg false discovery rate correction) are reported for the pairwise comparisons T vs TG3KO, T vs WT, T vs G3KO, TG3KO vs WT, TG3KO vs G3KO, and WT vs G3KO. **B–D)** present the results of Gene Set Enrichment Analysis (GSEA) performed against the REACTOME pathway database for the comparisons T vs TG3KO (**B**), T vs WT (**C**), and TG3KO vs WT (**D**). For each enriched pathway, the normalized enrichment score (NES), nominal *p* value, adjusted *p* value, and pathway description are provided. **E)** provides a summary of significantly enriched Reactome pathways identified in the T vs TG3KO, T vs WT and TG3KO vs WT comparisons, indicating for each pathway the direction of enrichment (positive NES reflecting over-enrichment in the first group of the comparison and negative NES indicating under-enrichment).

**Suppl. Table S2. Differential protein expression and pathway enrichment analyses of entorhinal cortex proteomic data**. This table comprises five sheets (A–E). **A)** contains the results of differential expression analysis performed using the *limma* framework on hippocampal proteomic data, including proteins that passed quality control and filtering criteria. For each protein, log2 fold change (logFC), raw *p* value, and multiple-testing adjusted *p* value (adjusted.p.value; Benjamini–Hochberg false discovery rate correction) are reported for the pairwise comparisons T vs TG3KO, T vs WT, T vs G3KO, TG3KO vs WT, TG3KO vs G3KO, and WT vs G3KO. **B–D)** present the results of Gene Set Enrichment Analysis (GSEA) performed against the REACTOME pathway database for the comparisons T vs TG3KO (**B**), T vs WT (**C**), and TG3KO vs WT (**D**). For each enriched pathway, the normalized enrichment score (NES), nominal *p* value, adjusted *p* value, and pathway description are provided. **E)** provides a summary of significantly enriched Reactome pathways identified in the T vs TG3KO, T vs WT and TG3KO vs WT comparisons, indicating for each pathway the direction of enrichment (positive NES reflecting over-enrichment in the first group of the comparison and negative NES indicating under-enrichment).

**Suppl. Table S3. Gene set enrichment analysis (GSEA) of KEGG pathways in hippocampus and entorhinal cortex proteomic datasets.** This table comprises six sheets (A–F), all reporting the results of GSEA performed against the KEGG pathway database. **A–C)** correspond to hippocampal proteomic data and present enrichment results for the comparisons T vs TG3KO (A), T vs WT (B), and TG3KO vs WT (C). **D–F)** correspond to entorhinal cortex proteomic data and present the same comparisons in the same order: T vs TG3KO (**D**), T vs WT (**E**), and TG3KO vs WT (**F**). For each enriched KEGG pathway, the normalized enrichment score (NES), nominal p value, and multiple-testing adjusted *p* value are provided, together with the pathway description.

**Suppl. Table S4. Differential expression and pathway enrichment analyses of white matter proteomic data.** This table comprises nine sheets (A–I). **A)** contains the results of differential expression analysis performed using the *limma* framework on white matter proteomic data, including proteins that passed quality control and filtering criteria. For each protein, log2 fold change (logFC), raw *p* value, and multiple-testing adjusted *p* value (adj.p.Val; Benjamini–Hochberg false discovery rate correction) are reported for the pairwise comparisons T vs TG3KO, T vs WT, T vs G3KO, TG3KO vs WT, TG3KO vs G3KO, and WT vs G3KO. **B–D)** Present the results of GSEA performed against the Reactome pathway database for the comparisons T vs TG3KO (B), T vs WT (C), and TG3KO vs WT (D). For each enriched pathway, the normalized enrichment score (NES), nominal *p* value, adjusted *p* value, and pathway description are provided. **E)** provides a summary of significantly enriched Reactome pathways identified in at least the T vs TG3KO and T vs WT comparisons, indicating for each pathway the direction of enrichment (positive NES reflecting over-enrichment in the first group of the comparison and negative NES indicating under-enrichment) together with its enrichment status in the TG3KO vs WT comparison. **F–H)** report the results of GSEA performed against the KEGG pathway database for the comparisons T vs TG3KO (F), T vs WT (G), and TG3KO vs WT (H), including the normalized enrichment score (NES), nominal *p* value, adjusted *p* value, and pathway description. **I)** Summarizes the significantly enriched KEGG pathways across comparisons, indicating the direction of enrichment and their comparative status between genotypes.

**Antibody list**

| Antibody | Host | Concentration | Supplier | Catalog; RRID |
| --- | --- | --- | --- | --- |
| Phospho-tau 202, 205; AT8 | Mouse | 1:500 | Thermo Scientific | MN1020; AB_223647 |
| MC1 | Mouse | 1:500 | Peter Davies lab | N/A; AB_2314773 |
| Iba1 | Rabbit | 1:500 | Fujifilm Wako | 019-19741; AB_839504 |
| Galectin-3 | Goat | 1:1000 | R&D | AF1197; AB_2234687 |
| Myelin Basic Protein | Rat | 1:1000 | Millipore | MAB386; AB_94975 |
| Synaptophysin | Rabbit | 1:250 | Abcam | ab14692; AB_301417 |
| PSD-95 | Mouse | 1:250 | Millipore | MAB1596; AB_2092365 |
| Olig2 | Goat | 1:250 | R&D | AF2418; AB_2157554 |
| CC1 (APC) | Mouse | 1:200 | Millipore | OP80; AB_2057371 |
| CLEC7A (Dectin-1) | Rat | 1:250 | Invivogen | mabg-mdect; AB_2753143 |
| Axl | Goat | 1:500 | R&D | AF854; AB_355663 |
| MAP2 | Chicken | 1:1000 | Abcam | ab92434; AB_2138147 |
| Phospho-tau 396, 404; PHF-1 | Mouse | 1:500 | Peter Davies lab | N/A; AB_2315150 |
| NDUFB8 | Mouse | 1:100 | Abcam | ab110242; AB_10859122 |
| Parvalbumin | Rabbit | 1:500 | Swant | PV25; AB_10000344 |
